# LHR and Gαs trafficking drive sustained cAMP signalling from endosomes to control steroidogenesis

**DOI:** 10.1101/2025.04.30.651471

**Authors:** Juliette Gourdon, Maya Haj-Hassan, Vinesh Jugnarain, Christophe Gauthier, Sabine Alves, Florian Guillou, Éric Reiter, Frédéric Jean-Alphonse

## Abstract

A growing number of G protein-coupled receptors (GPCRs) signal through G proteins from intracellular compartments following endocytosis. While for few receptors the physiological relevance of such mechanism has been established, the relationships between the spatio-temporal organization of cellular signalling and the physiological responses are still to be elucidated for most receptors. Signalling by the luteinizing hormone receptor (LHR) is essential to regulate sex steroids production in gonads, but how G protein-dependent signals from endosomes are functionally important for steroidogenesis remains unexplored. Here, we demonstrate that transient LHR activation promotes a prolonged Gαs/cAMP signalling from endosomes, which requires ligand-induced independent receptor and Gαs trafficking. We show that endosomal trafficking is specifically required for ligand-induced cAMP accumulation in the nucleus and gene expression, as well as for steroids production in Leydig cells. Thus, this study contributes to further understand the molecular mechanisms by which LHR, through signalling compartmentalization, controls gonadal steroidogenesis.

## 2. Introduction

G protein-coupled receptors (GPCRs) regulate multiple essential physiological processes, and abnormalities in their signalling often lead to pathological conditions^1,2^. A growing number of GPCRs are shown to signal through G proteins from intracellular sites following endocytosis. Intracellular signalling events have been described to occur from several cellular localizations, including the trans-Golgi network (TGN), the nuclear envelope, mitochondria, cilia, and endosomes. However, most GPCRs reported to engage G protein-dependent signalling after their internalization act at the early endosomes (EEs)^2–5^. Functional importance of G protein-dependent signalling from EEs has been described for some GPCRs, including calcitonin gene-related peptide (CGRP) receptor, parathyroid hormone receptor (PTHR), β2-adrenergic receptor (β2AR) or vasopressin type 2 receptor (V2R)^6–11^. While for few receptors the physiological relevance has been established, the impact of receptor trafficking and spatial and temporal organization of signalling on biological responses remains unknown for most of GPCRs. In addition, the role and mechanism of G protein trafficking in such endosomal signalling have been poorly studied. Notably, previous studies reported that G protein-dependent signalling of a given GPCR from different cellular compartments is associated with distinct cellular responses and physiological functions^8,11^. Therefore, deciphering the role of the different signalling compartments is of great importance to better understand physiological functions regulated by a GPCR and further develop specific therapies.

The luteinizing hormone receptor (LHR) is a class A GPCR, whose signalling regulates steroidogenesis and ovulation in gonads and is therefore essential for reproduction^12^. Upon stimulation by endogenous ligand, LHR internalizes in various endocytic vesicles with a main trafficking to atypical endosomes, called Very Early Endosomes (VEEs). While it is unclear whether the small fraction of receptors trafficking to EEs can signal from these compartments, the LHR has been shown to engage Gαs/cAMP signalling from VEEs^13–15^. VEEs are distinct from classical EEs by their smaller size and the lack of EEs’ typical markers, such as PI3P or EEA1^13^, but their biochemical characterization remains very limited. Therefore, studying VEEs-localized signalling is a challenging task, and it is complex to discriminate signalling arising from distinct endosomes populations. Nonetheless, the functional relevance of the whole LHR endosomal signalling remains to be determined. It has been proposed that LHR endosomal cAMP is necessary to oocyte meiosis resumption that precedes ovulation^16^. The main function of LHR expression in gonadal cells (i.e. Leydig cells in male, granulosa and theca cells in female) is to produce steroids necessary to support gonads’ activity. However, the impact of LHR signalling from endosomes in the regulation of steroidogenesis has not been investigated. So far, engagement of endosomal Gαs/cAMP signalling by LHR has been demonstrated using LH stimulation^13,14,16^. The human receptor possesses a second endogenous ligand, the human chorionic gonadotropin (hCG), which is broadly used for assisted reproduction treatments (ART)^12^. Whether hCG can engage endosomal Gαs/cAMP signalling downstream LHR has not been established yet.

In this study, we investigate the physiological relevance of both LHR and Gαs trafficking for the induction of steroidogenesis. We first show that hCG engages sustained cAMP signalling from endosomes in a similar manner than LH. We demonstrate that LHR endocytosis is required to promote Gαs activation, cAMP signalling and cAMP responsive element (CRE)-dependent gene expression. We show that LHR-induced Gαs translocation from the plasma membrane (PM) to the cytosol and endosomes is independent of LHR endocytosis, and is necessary for LHR sustained cAMP signalling from endosomes. Finally, using both cell lines and mouse primary Leydig cells, we demonstrate that LHR and Gαs endocytosis are essential for steroids production. Therefore, this study provides the first evidence that LHR spatially organizes Gαs activity, and that cAMP signalling from endosomes is essential to support steroidogenesis.

## 3. Results

### 3.1. LHR stimulation generates a prolonged Gs activation and a sustained cAMP signalling

A growing number of GPCRs have been shown to elicit Gαs/cAMP signalling from endosomes^2,17^. However, the intensity and duration of this endocytic signalling varies from receptor to receptor, is dependent on the ligand’s pharmacological properties, and is correlated to the receptor’s dynamic association with β-arrestins^17^. We compared the signalling dynamics of LHR with those of two GPCRs well-known to promote endosomal Gαs/cAMP, namely, the β2AR and the V2R. The β2AR displays acute but transient endosomal signalling, while the V2R stimulated by either arginine-vasopressin (AVP) or oxytocin (OT) has prolonged or short-lived cAMP signalling, respectively^8,18,19^. To ascertain the type of endosomal cAMP signalling triggered by the LHR, we first compared the cAMP signalling dynamics of the β2AR and the V2R in HEK293A cells stimulated with various concentrations of isoproterenol (ISO) or AVP, respectively, using the cAMP Bioluminescence Resonance Energy Transfer (BRET) sensor NLuc-Epac-VV^20^. Continuous stimulation of β2AR or V2R with various ligand doses showed saturating cAMP signals for both receptors over 60 minutes acquisition time, even at low nanomolar range of ligand concentrations (figure 1A, 1C). This did not reveal the transient cAMP signalling profile of β2AR, likely due to continuous stimulation of cell surface receptors not yet internalized and/or to receptors recycled back to the PM and re-sensitized. Therefore, continuous stimulation did not allow to assess both the activation and desensitization processes. To avoid this limitation and to better discriminate cAMP signalling dynamics, we performed “pulse-chase” stimulation for 60 seconds, followed by the ligand washout. This approach allowed us to monitor both the intensity and duration of the signal as well as the receptor and phosphodiesterases (PDEs)-mediated desensitization phase. As expected, “pulse-chase” stimulation of the β2AR revealed a two-phase cAMP response, with a rapid and transient cAMP peak followed by a reduced but prolonged cAMP signalling phase (figure 1B). For the V2R, transient stimulation with AVP showed a prolonged cAMP production with a much slower decay of the sustained cAMP signal (figure 1D). As for β2AR and V2R, continuous stimulation of the LHR with either LH or hCG led to saturated and sustained cAMP signals (figure 1E, S1A). In the “pulse-chase” stimulation assay, LH and hCG both promoted a prolonged cAMP signalling with very limited desensitization over the 60 minutes acquisition (figure 1F, S1B) and with comparable efficacies (figure 2G). These profiles resembled to the AVP-stimulated V2R, suggesting that LHR may signal from endosomes in a V2R-like manner. Interestingly, whereas “pulse-chase” stimulation with the highest ligand concentration importantly reduced the maximal cAMP level detected downstream β2AR and V2R compared to continuous stimulation, it did not significantly change the maximal cAMP level detected downstream LHR (figures 1A-F, S1). To assess whether sustained cAMP dynamics by LHR relies on Gs activation or decreased PDEs activity, we used the ONE-GO Gs biosensor to monitor the Gs-GTP dynamics^21^. Following the stimulation of LHR with 10 nM hCG in a continuous or transient manner, we observed a rapid Gs activation followed by a slow decrease (figure 1G, 1H). These results suggest that LHR activates Gs in a sustained manner and undergoes a slow rate of desensitization. However, with 1 nM hCG, we observed a remarkable difference between continuous and transient stimulation in the potency to activate Gs (figure 1G, 1H), comparable to what we observed with cAMP measurements (figure 1E, 1F). Our observations suggest that the long-lasting cAMP signalling by the LHR is associated with a prolonged Gs activation.

**Figure 1:**
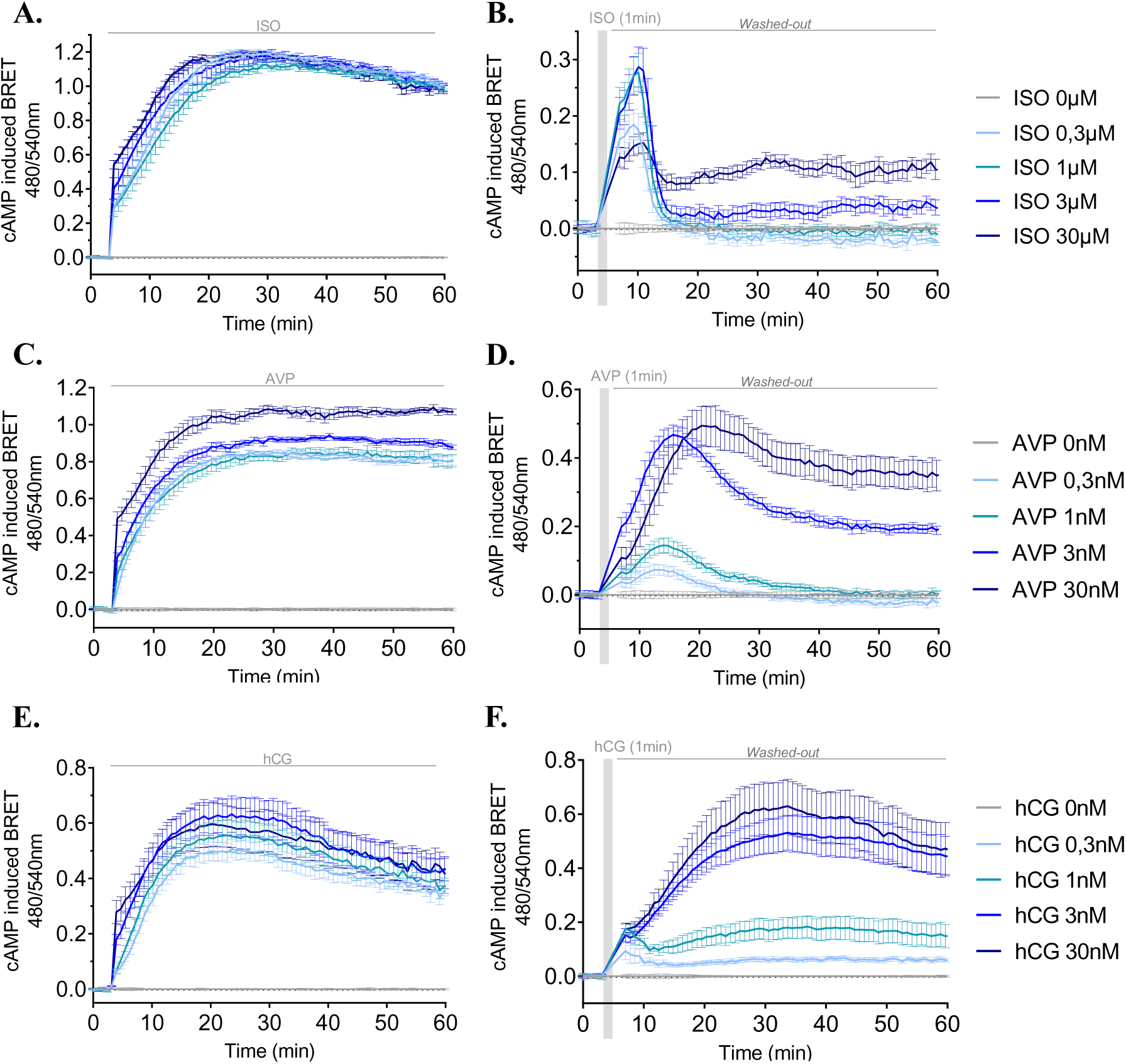
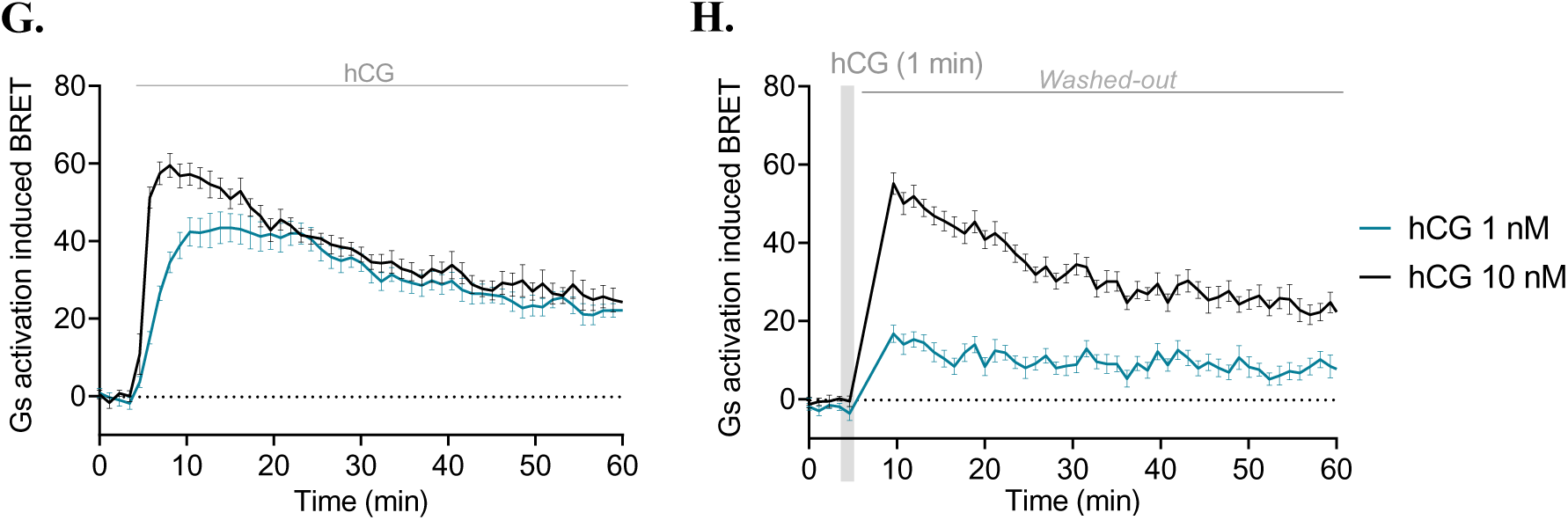
LHR stimulation generates a prolonged Gs activation and a sustained cAMP signalling. **(A, B)** β2AR cAMP signalling induced by continuous (A) or 60-seconds (B) isoproterenol (ISO) stimulation. **(C, D)** V2R cAMP signalling induced by continuous (C) or 60-seconds (D) arginine-vasopressin (AVP) stimulation. **(E, F)** LHR cAMP signalling induced by continuous (E) or 60-second (F) hCG stimulation. **(G, H)** ONE-GO Gs sensor showing the activation of Gs induced by continuous (G) or 60-seconds (H) hCG stimulation. For all graphs, **N=3 (n=9)**, means ± SEM.

**Figure 2:**
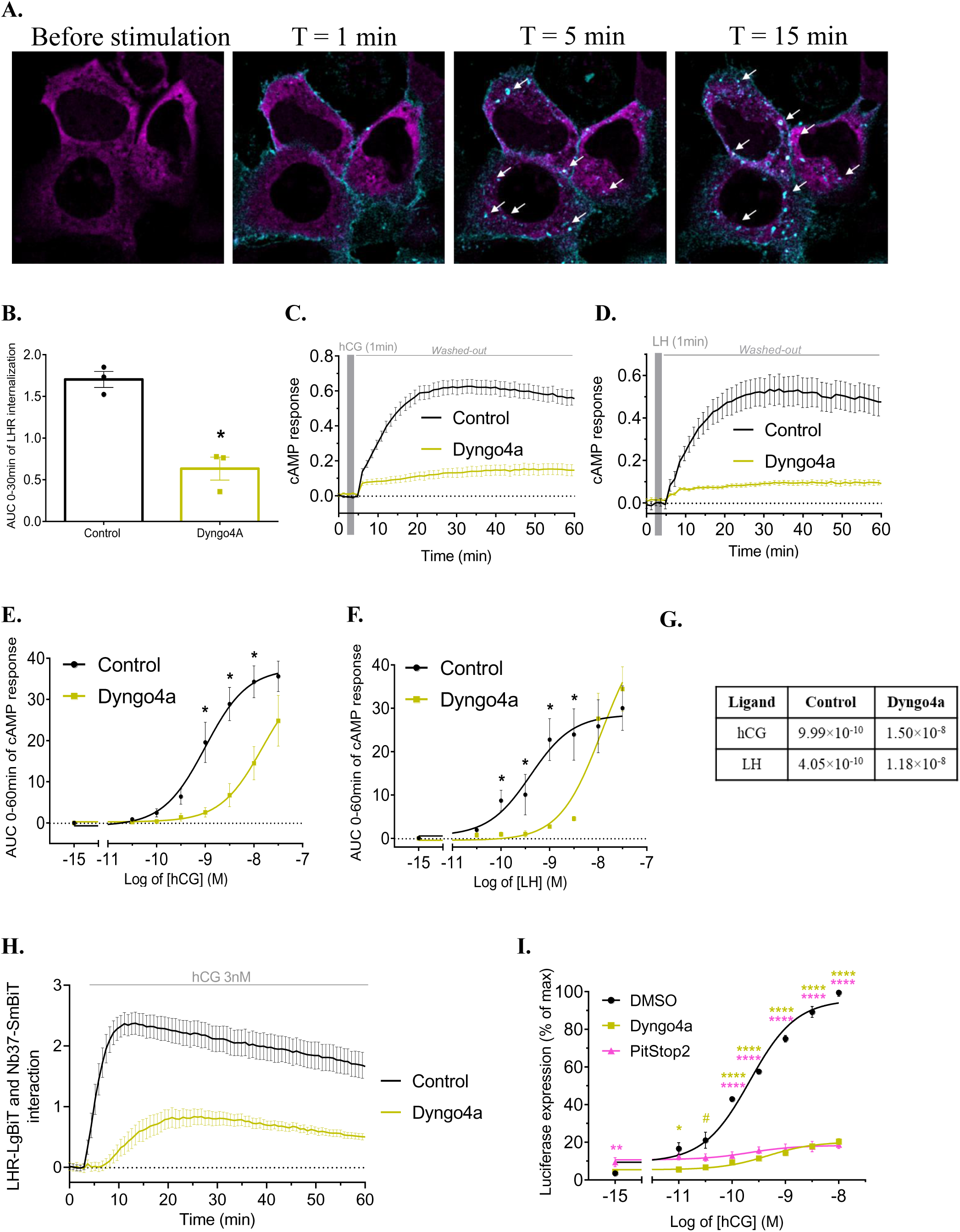
LHR endocytosis is essential to its signalling dynamics. **(A)** HEK293A cells transiently expressing LHR and treated with 100 nM hCG-mNG (cyan) during 60 seconds before ligand washout. At times 5 min and 15 min, arrows show colocalization between hCG-mNG positive endosomes and APPL1-mCherry (magenta). **(B)** Bystander BRET assay measuring hCG-induced LHR-RLuc8 endocytosis in HEK293A cells. Area under the curve (AUC) of LHR endocytosis between 0 and 30 minutes of continuous 30 nM hCG stimulation, with or without 30µM Dyngo4a. **N=3** experiments performed in triplicates, means ± SEM. **(C, D)** BRET measurements of time courses of cAMP using the biosensor NLuc-Epac-VV. LHR was stimulated either by 3 nM hCG (C) or 3 nM LH (D) for 60 seconds, with or without 30µM Dyngo4a. **N=3 (n=9)**, means ± SEM. **(E, F)** Dose-responses curves of the AUC between 0 and 60 minutes of cAMP production stimulated by hCG (E) or LH (F), with or without 30µM Dyngo4a. **N=3 (n=9)**, means ± SEM. **(G)** EC50 of cAMP response induced by 60-second stimulation with hCG or LH, calculated based on the 0-60 minutes AUC dose-responses. EC50 were calculated on data from three independent experiments performed in triplicates, and are indicated in molar. **(H)** Luminescence complementation assay monitoring the activation of Gs and the formation of a complex with LHR. Increased luminescence represents the recruitment of Nb37-SmBiT to the LHR-LgBiT by activated endogenous Gαs. **N=3 (n=9)**, means ± SEM. **(I)** CRE-driven luciferase gene expression after 6 hours continuous stimulation of HEK293A cells expressing LHR with different hCG concentrations, in presence of DMSO (control), 30µM Dyngo4a or 30µM PitStop2. Data were expressed in percentage of the control’s maximum. **N=3 (n=9)**, means ± SEM. Comparisons to DMSO for a given condition are indicated with corresponding colour stars and hashes. # p-value = 0.068.

### 3.2. LHR endocytosis is essential to its signalling dynamics

To determine whether the hCG-induced sustained cAMP signalling was indeed due to cAMP production from endosomes, we assessed the role of LHR trafficking in this response. We designed a recombinant hCG fused to the bright fluorescent protein mNeonGreen (hCG-mNG). First, we showed in HEK293A cells expressing LHR that transient hCG-mNG stimulation led to its accumulation in endosomes, with partial colocalization with the VEEs marker APPL1 (figure 2A). We further confirmed LHR endocytosis using a bystander BRET approach, and showed that receptor endocytosis was mediated by a dynamin-dependent mechanism. Indeed, treatment with Dyngo4a, a dynamin GTPase inhibitor^22^, significantly reduced LHR internalization (figure 2B). Inhibiting dynamin-dependent endocytosis strongly impaired hCG-induced cAMP production, significantly reducing global cytosolic cAMP levels within 60 minutes (figure 2C, 2E). Dyngo4a treatment affected LH-induced cAMP response in a similar manner (figure 2D, 2F), confirming the importance of post-endocytic signalling in the whole cellular cAMP response. Dose-responses analysis showed that blocking the dynamin-dependent endocytosis dramatically reduced the EC50 of cAMP production downstream LHR stimulated by either LH or hCG (figure 2G). The clathrin inhibitor PitStop2^23^ also strongly impaired cAMP production, further confirming the importance of receptor endocytosis in cAMP signalling (figure S2). Then, we used a nanoluciferase complementation strategy to monitor the formation of a complex between LHR and active Gαs. Upon agonist stimulation, the LHR C-terminally fused to the LgBiT nanoluciferase portion (LHR-LgBiT) and the Gs sensor Nb37^18^ fused to the SmBiT nanoluciferase portion (Nb37-SmBiT) showed rapid association, followed by a slow decrease (figure 2H). Interestingly, Dyngo4a treatment delayed and strongly impaired the association of Nb37-SmBiT with LHR-LgBiT (figure 2H), suggesting that the receptor did no longer form a complex with activated Gαs. We next assessed the consequences of treatment with either Dyngo4a or PitStop2 on CRE-dependent luciferase reporter gene expression, and revealed that inhibition of LHR endocytosis led to almost complete inhibition of hCG-induced gene expression (figure 2I). Altogether, this demonstrated that the majority of the LHR Gαs-cAMP response occurs after receptor endocytosis, and that signalling from endosomes is essential to control CRE-dependent gene expression.

### 3.3. Ligand-induced endocytosis of Gαs is required for its activation and for sustained cAMP signalling

The mechanism and role of Gαs trafficking in GPCR signalling from endosomes remain poorly understood. Although endosomal localization of active Gαs is necessary for cAMP signalling, it is still unclear whether all GPCRs stimulating endosomal signalling activate G protein heterotrimers already present in endosomes, or recruit Gs proteins in these compartments for local activation^24,25^. Using a confocal microscopy approach, we assessed Gαs localization and trafficking using a Gαs fused to yellow fluorescent protein (Gαs-YFP). Before stimulation, Gαs-YFP was mainly localized at the PM, but progressively dissociated from it upon LHR stimulation (figure 3A). BRET experiments further confirmed this hCG-induced Gαs endocytosis (figure 3B). Following PM dissociation, Gαs-YFP first accumulated in the cytosol, and was then observed in endocytic vesicles, as shown by partial colocalization with the VEEs marker APPL1 and the EEs marker Rab5 (figure 3C, 3D). Interestingly, this traffic was not affected by Dyngo4a treatment (figures 3B, S3), suggesting that Gαs endocytosis was dynamin-independent, and that Gαs could traffic independently of the receptor upon agonist stimulation. To investigate the role of Gαs trafficking in endosomal signalling, we generated a Gs variant harbouring a N-terminal myristoylation site (Myr-Gs), to stabilize Gαs localization at the PM and limit its release from it^53^. BRET assays and confocal imaging showed that upon hCG stimulation, Myr-Gs mostly remained at the PM (figures 3E, S4A). We then assessed the consequences of decreased Gαs trafficking to endosomes on cAMP dynamics. HEK293A cells with CRISPR/Cas9 knock-out for Gαs (HEK293ΔGs) were transfected with either WT-Gs or the Myr-Gs. Though LHR pulse-chase stimulation with hCG induced the increase of cAMP production in both conditions, Myr-Gs had a significantly reduced capacity to promote sustained cAMP signalling (figure 3F, 3G). This decrease in sustained cAMP signalling was consistent with a reduced recruitment of Nb37-SmBiT to LHR-LgBiT (figure S4B, S4C). Overall, these results demonstrated that Gαs requires LHR activation for its trafficking away from the PM, independent of the receptor’s endocytic route. Moreover, they showed that Gαs redistribution to endosomal membranes plays an essential role for LHR endosomal signalling.

**Figure 3:**
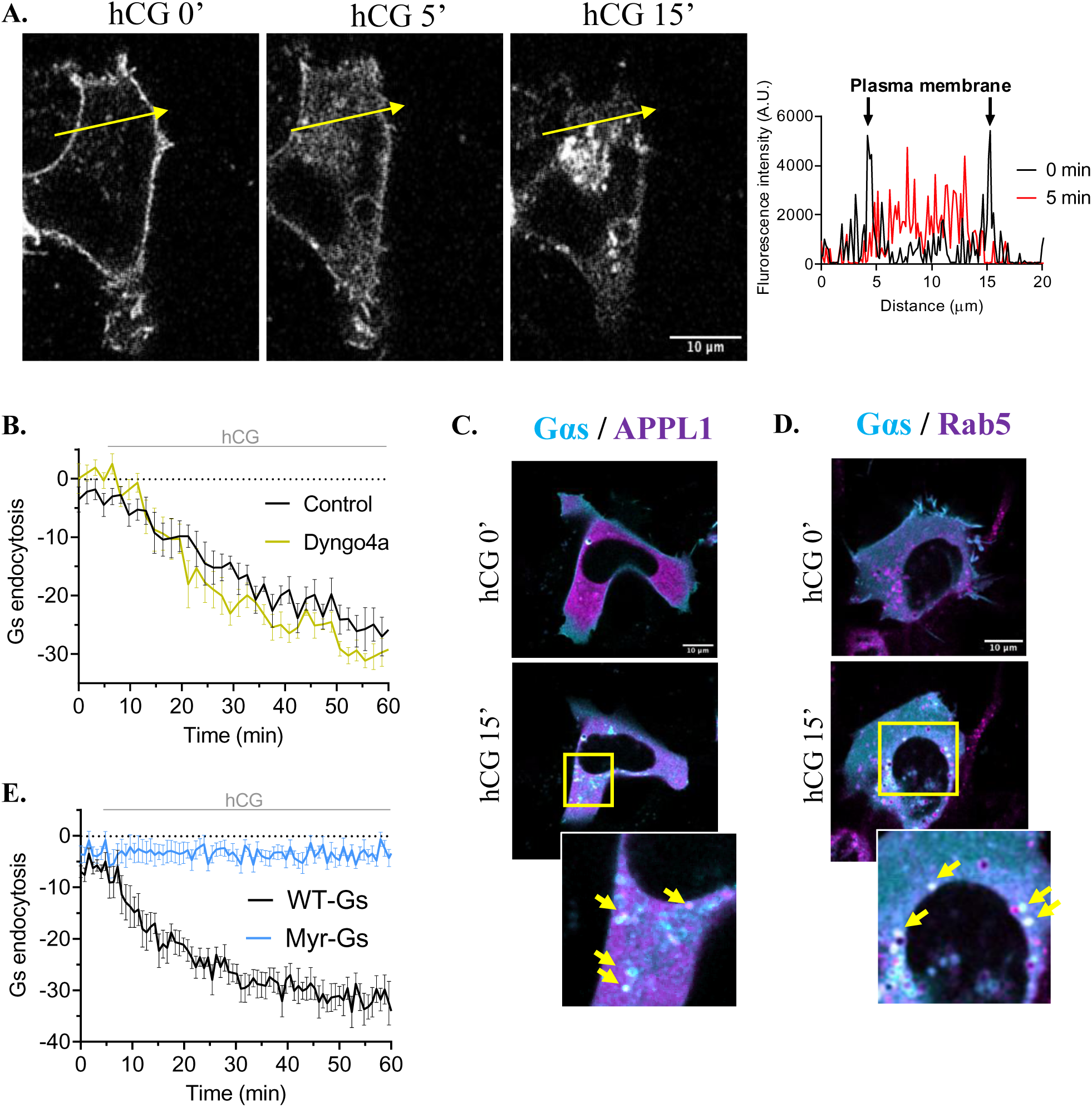
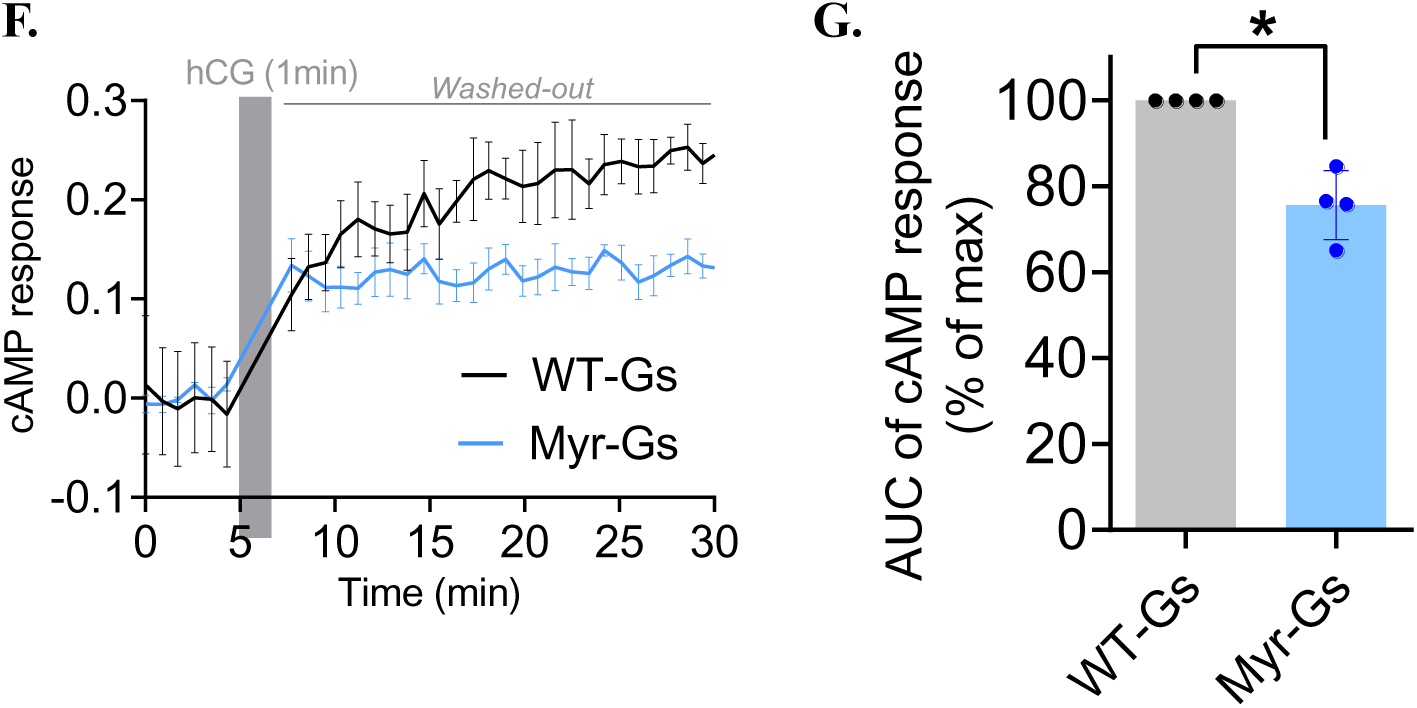
Ligand-induced endocytosis of Gαs is required for its activation and for sustained cAMP signalling. **(A)** Gαs-YFP cellular localization over time after 30 nM hCG stimulation of HEK293A cells transiently expressing LHR. Representative pictures of Gαs-YFP cellular localization without ligand stimulation (left) and after 5 minutes (middle) or 15 minutes (right) hCG stimulation. Scale bar = 10 µm. Fluorescence intensity across cell width was quantified before (0min, in black) and after 5 minutes (5min, in red) hCG stimulation. **(B)** BRET assay measuring the dissociation of YFP-Gαs from the plasma membrane, upon 30 nM hCG stimulation with or without of 30µM Dyngo4a. **N=3 (n=9)**, means ± SEM. **(C, D)** Representative images from live cell imaging showing colocalization of YFP-Gαs (cyan) with either APPL1-mCherry (C) or Rab5-mScarlet-I (D) (magenta), before (top) or after 15 minutes (bottom) 30 nM hCG stimulation. Yellow arrows in insets (bottom right) show endosomes positive for both markers. Scale bar = 10 µm. **(E)** BRET assay comparing the endocytosis of the WT Gαs-YFP and the Myr-Gαs-YFP variant in HEK293ΔGs cells, following 30 nM hCG stimulation. **N=3 (n=9)**, means ± SEM. **(F)** BRET assay comparing the cAMP kinetics profiles of HEK293ΔGs cells transiently expressing LHR and co-expressing either WT-Gαs or Myr-Gαs, stimulated with 0.3nM hCG during 60 seconds. **N=1 (n=3) representative from N=3 independent experiments**, means ± SEM. **(G)** Quantification of the area under the curve from 0-30 minutes cAMP response. **N=3**, means ± SEM.

### 3.4. Plasma membrane-localized Gαs inhibition prevents Gαs translocation, cytosolic and nuclear cAMP accumulation, and gene transcription

To go further into deciphering the contribution of ligand-induced Gαs activation and translocation, and since there are no commercially available Gαs inhibitors, we used the Nb37 biosensor, that was initially developed to detect Gαs activation^18^. The Nb37 recognizes and stabilizes the nucleotide-free state intermediate activation conformation of Gαs, thus blocking guanosine triphosphate (GTP) incorporation and Gαs function. We aimed to determine whether we could use Nb37 as a competitor for both GTP and Gβγ, and thus as a dominant negative tool to block cAMP signalling. We fused the Nb37 to a PM-addressing sequence (Nb37-CAAX), and showed that its expression prevented hCG-induced Gαs dissociation from the PM (figure 4A-C). To assess the dominant-negative effect of Nb37-CAAX on Gs activation, we monitored Gαs-Gβγ interaction using the SmBiT-LgBiT complementation assay^28^. As expected, hCG induced a reduction in the luminescence, due to G protein activation and Gαs dissociation from Gβγ (figure S5A). Next, we compared the effect of overexpression of either the nucleus-targeted Nb37 (Nb37-3xNLS) or Nb37-CAAX on Gαs-Gβγ interaction in a dose-dependent manner. Our results showed that only Nb37-CAAX interfered in the heterotrimeric complex, in both basal and stimulated conditions (figures 4D, S5B). Recent studies suggested that GPCRs localization in endosomes could allow better cAMP diffusion into the nucleus to promote gene expression^26^. To assess the role of Gαs trafficking and activity in these functions, we generated a new cAMP BRET sensor, based on the FRET sensor CUTie developed to maintain similar properties whatever its cellular localization^27^. We fused our NanoCUTie BRET sensor either to a nucleus exclusion signal (NES) for cytosolic localization, or to a nuclear-localization sequence (NLS) (figure S6). Because of the sensor’s sensitivity, we could not perform “pulse-chase” ligand stimulation. Upon modest overexpression of Nb37-CAAX (3ng plasmid), hCG-induced cytosolic cAMP accumulation was only partially reduced. In contrast, nuclear cAMP accumulation was completely abolished (figure 4E-G). These results suggested that ligand-induced Gαs translocation to endosomes contributes to the whole cAMP response but is specifically required for cAMP accumulation at the nucleus. Consistent with these observations and with the literature^26^, upon 6 hours hCG stimulation, Nb37-CAAX overexpression significantly inhibited ligand-induced CRE-dependent luciferase reporter gene expression (figure 4H). Interestingly, in this latter assay Nb37-CAAX was expressed in high quantity (30ng plasmid), and the inhibition of gene expression was only partial. It is to be noted that nuclear cAMP levels were monitored within the first hour following receptor stimulation, whereas gene expression level was measured after 6 hours of agonist stimulation. Therefore, it is possible that within 6 hours, some cytosolic cAMP could diffuse to the nucleus and stimulate gene expression. When considering that LHR endocytosis inhibition led to full inhibition of CRE-dependent gene expression (figure 2I), it could also be assumed that LHR endocytosis would primarily drive hCG-induced gene expression through both Gαs and other signalling partners.

**Figure 4:**
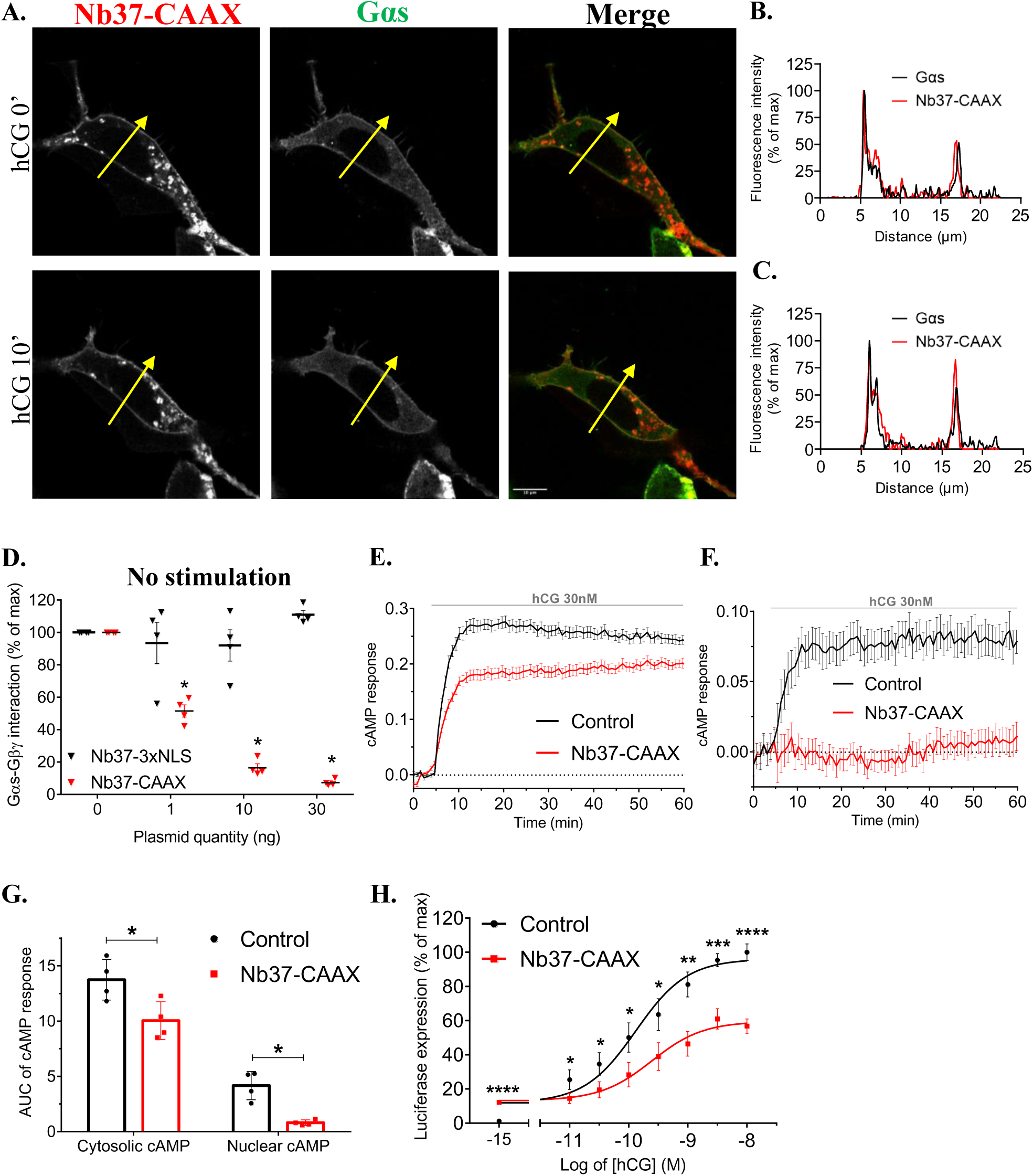
Plasma membrane-localized Gαs inhibition prevents Gαs translocation, cytosolic and nuclear cAMP accumulation, and gene transcription. **(A)** Cellular localization of Gαs-YFP and Nb37-CAAX-mScarlet in absence (top) or after 10 minutes (bottom) of 30 nM hCG stimulation. Scale bar = 10 µm. **(B, C)** Fluorescence intensities from (A) for Gαs-YFP (in black) and Nb37-CAAX-mScarlet (in red) across cell width, in absence (B) or after 10 minutes (C) of hCG stimulation. **(D)** Luminescence complementation assay monitoring the Gαs and Gβγ ligand-induced dissociation in presence of different amounts of transfected Nb37-NLS (nucleus localized) or Nb37-CAAX (plasma membrane localized). AUC of Gαs-LgBiT and Gβ1-SmBiT interaction between 0 and 10 minutes, in absence of ligand. Data were expressed as the percentage of control’s maximum. **N=4** independent experiments performed in triplicates, means ± SEM. **(E, F)** cAMP time courses induced by continuous 30 nM hCG stimulation in cells expressing either the cytosolic cAMP biosensor NanoCUTie-NES (E) or the nuclear localized sensor NanoCUTie-NLS (F), and co-expressing 3ng plasmid/well of mCherry (control) or Nb37-CAAX. **N=4 (n=12)**, means ± SEM. **(G)** AUC from cytosolic and nuclear cAMP responses measured between 0 and 60 minutes of hCG stimulation. **N=4**, means ± SEM. **(H)** CRE-driven luciferase gene expression after 6 hours hCG stimulation of HEK293A cells expressing LHR together with mCherry (control) or Nb37-CAAX-mScarlet (30ng plasmid/well). Data were expressed in percentage of the control’s maximum. **N=3 (n=9)**, means ± SEM.

### 3.5. Steroidogenesis in mouse Leydig cells requires LHR internalization and endosomal Gαs/cAMP signalling

Steroids production is a main physiological response mediated by LHR activity, and depends on ligand-induced Gαs/cAMP signalling and CRE-dependent gene transcription^29^. Therefore, we questioned the importance of both LHR and Gαs trafficking in the induction of steroidogenesis. To that purpose, we used mouse Leydig Tumour Cells-1 (mLTC-1), a cell line that retained endogenous expression of the mouse LHR (mLHR) and the ability to produce steroids^30^. We first showed that as for human LHR (hLHR) in HEK293A cells, mLHR stimulated with hCG-mNG in a “pulse-chase” manner underwent endocytosis (figure 5A). We next measured the cAMP dynamics in mLTC-1 cells using the recombinant hCG. As in HEK293A cells overexpressing hLHR, hCG induced a long-lasting cAMP response, that was strongly diminished when cells were treated with endocytosis-disrupting drugs Dyngo4a and PitStop2 (figures 5B, 5C, S7). All these results strongly suggested that our observations on hLHR in HEK293A cells were consistent with the behaviour of endogenous mLHR in gonads cells, thus validating the use of mLTC-1 cells to study the role of LHR endosomal signalling in steroidogenesis.

**Figure 5:**
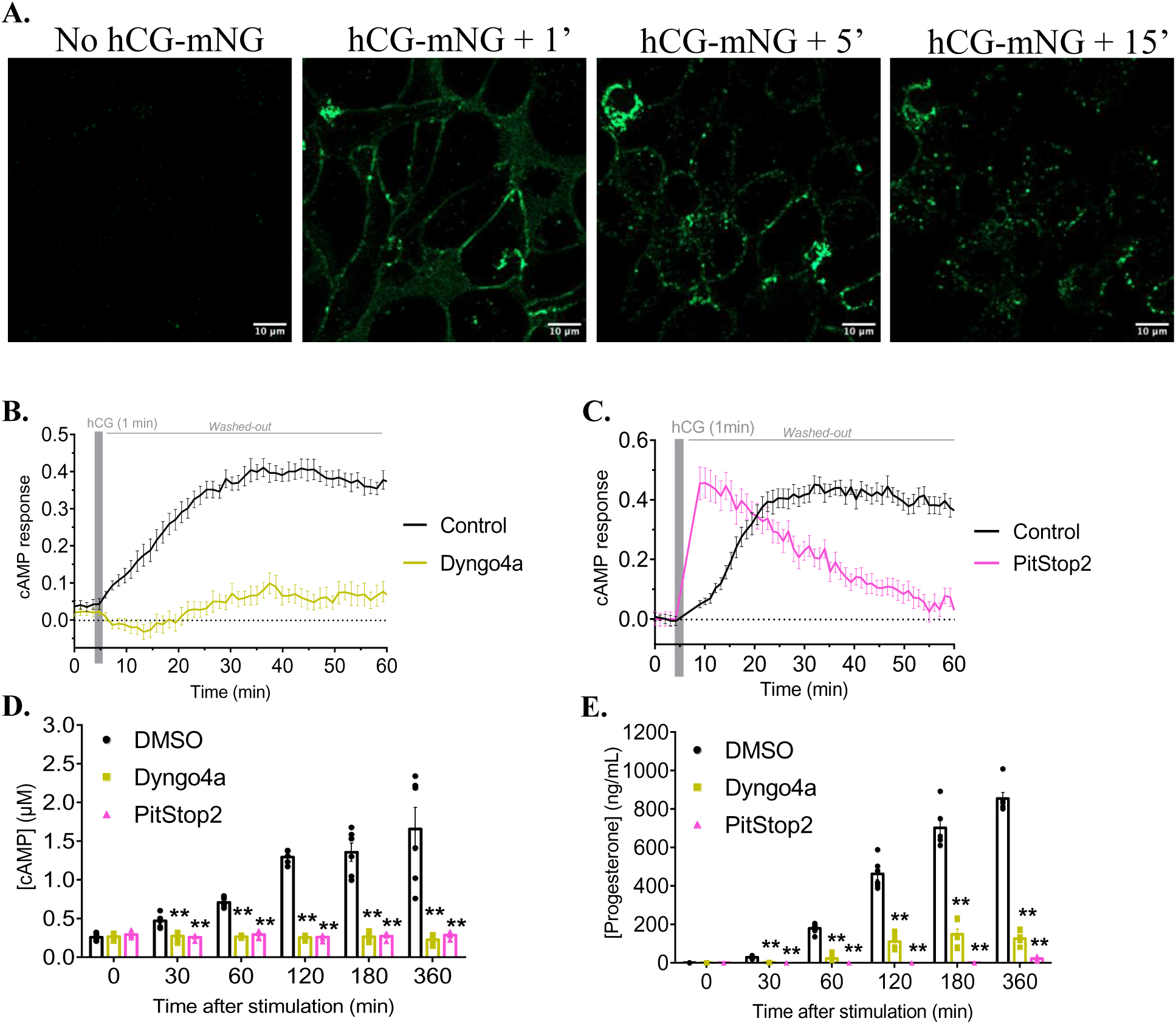
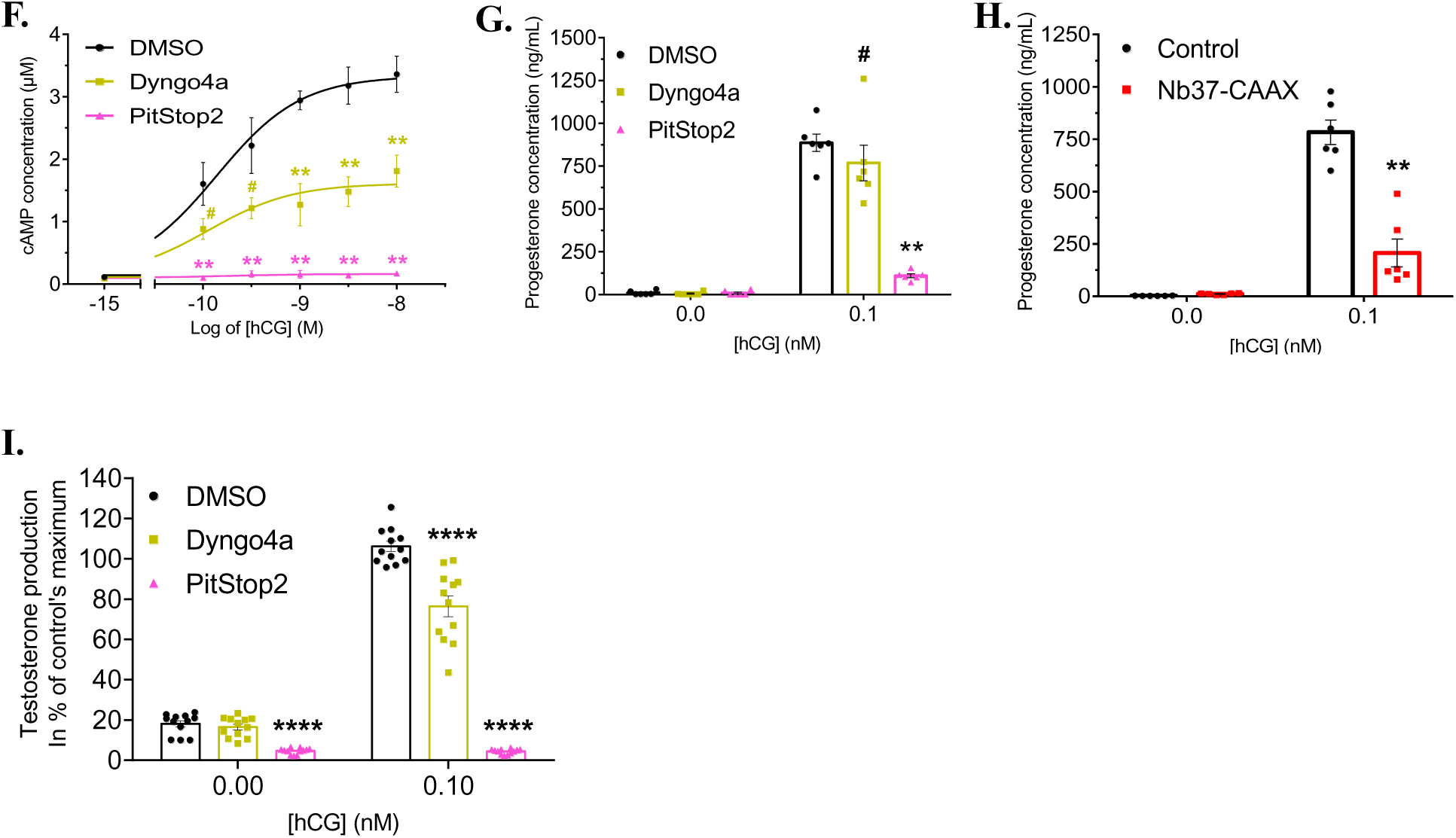
Steroidogenesis in mouse Leydig cells requires LHR internalization and endosomal Gαs/cAMP signalling. **(A)** Stimulation of mLTC-1 cells with 100 nM hCG-mNeonGreen for 120 seconds, before washout. Cellular localization of hCG-mNeonGreen before stimulation (no hCG-mNG), and 1, 5 or 15 minutes after stimulation. Scale bar = 10 µm. **(B, C)** Cytosolic kinetics of cAMP signalling in mLTC-1 cells, induced by 60-second 10nM hCG stimulation with or without 30µM Dyngo4a (B) or 30µM PitStop2 (C). **N=3 (n=9)**, means ± SEM. **(D, E)** Measurement of cAMP (D) and progesterone (E) concentrations in extracellular medium of mLTC-1 cells stimulated for 1 minute with 1nM hCG, in presence of DMSO (control), 30µM Dyngo4a or 30µM PitStop2. Supernatants were collected 30, 60, 120, 180 or 360 minutes after hCG stimulation. The 0 minutes condition corresponds to unstimulated cells for which supernatant was collected at 360 minutes. **N=3 (n=6)**, means ± SEM. **(F)** Dose-response curves of cAMP concentration in extracellular medium of mLTC-1 cells continuously stimulated for 3 hours with different hCG doses, in presence of DMSO, 30µM Dyngo4A or 30µM PitStop2. **N=3 (n=6)**, means ± SEM. Comparisons to DMSO for a given condition are indicated in corresponding colours. **(G)** Progesterone concentration in extracellular medium of mLTC-1 cells stimulated for 3 hours with 0.1nM hCG, in presence of DMSO, 30µM Dyngo4a or 30µM PitStop2. Progesterone dosages were performed on the same samples as cAMP dosages represented in (F). **N=3 (n=6)**, means ± SEM. # p-value = 0.066. **(H)** Progesterone concentration in extracellular medium of mLTC-1 cells transiently overexpressing mCherry (control) or Nb37-CAAX-mScarlet (80ng/well of 48-well plate, roughly equivalent to 30ng/well of 96-well plate). **N=3 (n=6)**, means ± SEM. **(I)** Measurement of testosterone concentration in extracellular medium of mouse primary Leydig cells stimulated for 3 hours with 0.1nM hCG, in presence of DMSO, 30µM Dyngo4a or 30µM PitStop2. Data were expressed as the percentage of control condition’s maximal response. **N=4 (n=12)**, means ± SEM.

Steroidogenesis is a relatively long-term cellular response to LHR stimulation, and we first questioned the temporality of its activation. It has been described that cAMP is extensively secreted, and that extracellular cAMP reflects the overall level of cAMP after 60 minutes of agonist stimulation^31^. We demonstrated that after only 60-second “pulse-chase” stimulation with 1nM hCG, cAMP progressively accumulated in mLTC-1 cells medium (figure 5D). The cAMP level almost reached its maximum 2 hours after ligand stimulation and remained protracted up to 6 hours. Dyngo4a and PitStop2 treatments abolished extracellular cAMP accumulation (figure 5D), underlining the indispensable role of LHR endocytosis in this response. mLTC-1 cells are able to produce steroids, but display a deficiency in the expression of the 17β-hydroxydehydrogenase isoform 3 (17βHSD3), resulting in accumulation of progesterone rather than testosterone^32^. Following “pulse-chase” hCG stimulation, progesterone was detected in mLTC-1 cells medium from 1 hour, and continued to progressively accumulate within 6 hours after agonist stimulation (figure 5E). When treating cells with Dyngo4a or PitStop2, progesterone accumulation over 6 hours was drastically inhibited (figure 5E). These data demonstrate for the first time that LHR endocytosis and endosomal cAMP signalling are crucial for steroids production. In addition, they suggest that steroidogenesis machinery in Leydig cells is already primed to rapidly respond to agonist stimulation, before amplification of the response by CRE-dependent expression of genes involved in steroidogenesis. Notably, we show that 60-second LHR stimulation is sufficient to induce steroids accumulation within hours.

We further wondered whether the importance of LHR endocytosis for cAMP and steroids accumulation would be the same under continuous agonist stimulation. Up to now, steroids production has usually been assessed after 3 hours of continuous ligand stimulation. In these conditions, hCG promoted a dose-dependent cAMP accumulation in mLTC-1 medium (figure 5F). At 1nM hCG, continuous ligand stimulation promoted twice higher cAMP accumulation than the one observed 3 hours after “pulse-chase” stimulation (figure 5D), likely due to re-engagement of receptors recycled back to the PM. Under 3 hours of continuous hCG stimulation, PitStop2 maintained a full inhibitory effect on cAMP accumulation, but Dyngo4a only had a partial inhibitory effect (figure 5F), suggesting this latter drug was permissive upon long-time incubation with mLTC-1 cells and allowed some internalization of recycled receptors. Interestingly, cAMP accumulated in presence of Dyngo4a was enough to stimulate progesterone production, though at a slightly lower level than in control condition, whereas PitStop2 treatment almost completely inhibited progesterone accumulation as expected (figure 5G). To confirm the role of Gαs in LHR endosomal signalling, we assessed hCG-induced progesterone production in mLTC-1 cells transiently expressing Nb37-CAAX. Compared to control, Nb37-CAAX expression engendered a strong inhibition of progesterone accumulation (figure 5H), suggesting Gαs activation and translocation to endosomes are necessary to support steroidogenesis in Leydig cells. Altogether, these results suggested that both LHR and Gαs endocytosis are essential to endosomal cAMP signalling and steroidogenesis. Finally, to confirm our observations in a more physiologically relevant model, we isolated Leydig cells from ten weeks-old mice. We performed a 3-hour continuous hCG stimulation and determined the concentration of testosterone accumulated in the primary Leydig cells medium. LHR agonist stimulation led to an important increase in testosterone concentration, which was partially or fully inhibited by Dyngo4a or PitStop2 treatment, respectively (figure 5I). Interestingly, PitStop2 treatment of primary Leydig cells reduced testosterone level even in absence of agonist stimulation (figure 5I), which was not observed in mLTC-1 cells. Primary Leydig cells were cultured and stimulated directly after extraction from mice testes. Therefore, it is possible that PitStop2, which was more efficacious than Dyngo4a during long-time incubation (figure 5F, 5G, 5I), inhibited endocytosis of receptors already bound to endogenous mouse LH, thus leading to a reduced testosterone production. Culture conditions did not allow to transfect primary cells with Nb37-CAAX, but the consistency of Dyngo4a and PitStop2 treatments effects between mLTC-1 and primary Leydig cells suggested the requirement of Gαs activation and translocation to endosomes would be conserved in primary cells. Altogether, our results demonstrate for the first time that both LHR and Gαs endocytosis are essential to drive sustained cAMP signalling from endosomes and control steroidogenesis in Leydig cells.

## 4. Discussion

GPCRs regulate many physiological functions, and their endocytic trafficking is critical to decode complex cellular signalling. Despite the growing number of GPCRs described to mediated G protein signalling from endosomes, very few studies reported the functional role of G protein-dependent signalling’s spatial organization for physiological or pathological responses. Previous studies reported that LHR, whose signalling regulates gonads activity and reproduction, mostly traffics to VEEs rather than classical EEs^13,14^. Despite this atypical trafficking, it has been suggested that LHR signalling from endosomes, as for other EEs-signalling Gs-coupled receptors, is critical to regulate specific functional responses to agonist stimulation. Indeed, LH-dependent endosomal localization has been shown to regulate ERK1/2 phosphorylation, and cAMP produced by internalized LHR is thought to regulate oocyte meiosis resumption that precedes ovulation^13,16^. In this study, we show that both endogenous ligands LH and hCG drive sustained LHR Gαs/cAMP signalling from endosomes. We demonstrate that LHR endosomal Gαs/cAMP signalling not only represents the greatest part of agonist-induced cAMP response, but specifically regulates steroidogenesis, one of the main physiological outcomes of LHR activity.

In an effort to study the role of post-endocytic cAMP, we first set up experimental conditions to properly assess cAMP dynamics, including production and degradation of the second messenger. Using the β2AR and the AVP-stimulated V2R as models of transient and prolonged cAMP signalling from endosomes, respectively^8,18,19^, we showed that “pulse-chase” ligand stimulation, but not continuous exposure to ligand, allowed to properly discriminate cAMP dynamics in terms of intensity and duration (figure 1A-D). By preventing continuous exposure to the ligand of cell surface-remaining receptors and potential stimulation of receptors recycled back to the PM, “pulse-chase” stimulation thus provided a powerful approach to decipher cAMP signalling arising from intracellular sites. LH and hCG were previously shown to display biased agonism at the LH receptor, hCG being more potent at triggering cAMP responses, β-arrestin recruitment to LHR, receptor endocytosis and steroid production^33–35^. In this study, we demonstrate that both LH and hCG stimulate sustained cAMP signalling downstream LHR, which is associated to acute and rather prolonged Gs activation (figures 1E-H, S1). Interestingly, we did not observe a greater potency (EC50) in inducing cAMP production for hCG compared to LH, but rather a greater efficacy (Emax) (figure 2E-G).

As inhibition of LHR endocytosis prevented sustained cAMP signalling, we confirmed that both endogenous ligands triggered cAMP production mainly from intracellular compartments (figures 2B-G, S2). Intriguingly, inhibition of dynamin or clathrin by Dyngo4a or PitStop2, respectively, did not affect hCG-induced cAMP dynamics in a similar manner. Whereas Dyngo4a treatment led to low level, slow cAMP accumulation, PitStop2 treatment promoted acute, rapid cAMP production followed by a rapid degradation and return to basal level following ligand washout (figures 2C, S2A). On one hand, this suggested cAMP accumulation almost exclusively originated from endosomes, whereas on the other hand it suggested LHR first mediated short, acute cAMP response at PM followed by sustained cAMP response from endosomes. However, as opposed to Dyngo4a, PitStop2 treatment shifted early LHR cAMP dynamics from progressive accumulation (≃ 10 minutes to reach the plateau) to rapid and acute response (≃ 5 minutes to reach the peak) (figure S2A). This led us to hypothesize that PitStop2 pre-incubation engendered massive accumulation of LHR at PM, and that the rapid cAMP increase would be due to stimulation of a large number of receptors at cell surface, unfollowed of sustained cAMP response because of the lack of receptor endocytosis and intracellular signalling. Nonetheless, both Dyngo4a and PitStop2 treatments demonstrated that LHR sustained cAMP signalling occurs from intracellular compartments. Interestingly, inhibition of LHR endocytosis impaired the formation of an LHR-active Gαs complex (figure 2H), suggesting that their coupling not only occurs at PM but also at intracellular sites. Furthermore, we show that intracellular sustained Gαs/cAMP signalling is essential to promote CRE-dependent gene expression (figure 2I). These results were consistent with previous observations on the β2AR, for which endosomal Gαs/cAMP signalling specifically regulates gene transcription^7,10^, and highlighted the functional importance of LHR post-endocytic signalling.

Activation of Gαs, which is required for cAMP production, is primarily known to occur at PM. However, studies have highlighted the requirement of active Gαs in endosomes to mediate cAMP signalling from these compartments. For the β2AR, it remains unclear whether Gs heterotrimers are already present in endosomes or recruited there by internalized GPCRs^24,25^. A recent study proposed that upon V2R stimulation, activated and depalmitoylated Gαs dissociates from PM, and that Gβγ internalized in endosomes in a receptor- and β-arrestin-dependent manner then stimulates Gαs translocation to endosomes^36^. Here, we demonstrate that LHR agonist stimulation induces Gαs translocation to APPL1-positive endosomes, in addition to other endocytic structures including Rab5-positive endosomes (figure 3A, 3C, 3D). Contrary to LHR endocytosis, Gαs translocation was not affected by Dyngo4a treatment (figures 3B, S3), showing that Gαs trafficking is independent of that of the receptor, consistently with recent findings on the V2R^36^. Nevertheless, inhibition of LHR endocytosis prevented Gαs activation (figure 2H), strongly suggesting that Gαs trafficking is essential to allow its optimal activation, mainly from intracellular sites. Accordingly, a myristoylated Gs variant, which displayed a stronger anchoring at the PM and a reduced capacity to translocate, had an impaired activation and capacity to promote cAMP signalling compared to the WT-Gs, further supporting the need for Gs’ release from the PM (figures 3E-G, S4). Gαs attachment to the PM utilizes both a palmitoylation and its interaction with Gβγ. Upon activation, Gαs proteins dissociate from Gβγ, and undergo rapid depalmitoylation by acyl protein thioesterases (APT) followed by release into the cytosol^37–40^. Interestingly, the mechanism of Gαs trafficking seems more complex, as APT pan-inhibitor was shown not to impact Gαs dissociation from the PM^25^. In addition, whether Gβγ dimer is required for LHR-dependent Gαs localisation to endosomes, as shown for the V2R or PTHR^36,41^, is still to be determined.

Taking advantage of the Nb37 properties^42^ in absence of available Gαs inhibitors, we deciphered further the contribution of ligand-induced Gαs activation and translocation to endosomes. We demonstrated that PM-addressed Nb37 (Nb37-CAAX), likely by scavenging Gαs, not only inhibited Gαs-Gβγ interaction in a dose-dependent manner, but also prevented ligand-induced Gαs translocation (figure 4A-C). Partial inhibition of Gαs activation and translocation by Nb37-CAAX reduced cytosolic cAMP response, and completely abolished cAMP accumulation at the nucleus within 60 minutes (figure 4E-G). We cannot completely exclude that reduction of cAMP response was due to a smaller number of Gαs available for coupling with LHR, rather than to Nb37-CAAX preventing Gαs translocation. However, effects of Nb37-CAAX on cytosolic cAMP response were consistent with our results with Myr-Gs, that remained fully available for coupling at the PM. Because of the limited sensitivity of our compartmentalized NanoCUTie BRET sensor, we had to assess compartment-specific cAMP dynamics under continuous ligand stimulation, which implies that cell surface-remaining receptors could continuously be stimulated. Even in these conditions, Nb37-CAAX expression, that reduced but not abolished cytosolic cAMP response, and so did not fully prevented Gαs activity, completely blocked cAMP accumulation at the nucleus (figure 4E-G). Altogether, this demonstrated that Gαs activity in endosomes is specifically required for LHR-dependent cAMP accumulation at the nucleus, as well as for the regulation of CRE-dependent gene expression (figure 4H). These results were consistent with a study on the thyroid-stimulating hormone receptor (TSHR), which demonstrated that Gαs/cAMP signalling from endosomes activates a third wave of cAMP signalling and CRE-dependent gene transcription at the nucleus^43^. More precisely, the authors reported that endocytic Gαs/cAMP signalling triggers calcium release from the endoplasmic reticulum, which can rapidly reach the nucleus and activate soluble adenylate cyclase (sAC) to promote cAMP production, and ultimately CRE-dependent gene expression. TSHR and LHR belong to the same family of glycoprotein hormones receptors. Whether they share a similar mechanism for cAMP nuclear accumulation, or LHR uses cAMP diffusion as suggested for other GPCRs like the β2AR^26^, would need further investigations. However, chemical inhibition of sAC in mLTC-1 cells was previously reported to negatively regulate LHR mediated cAMP production^44^. This suggests that LHR cAMP signalling could follow a three-waves model, involving cAMP production from PM and endosomes by Gαs, and from nucleus *via* sAC.

A major physiological outcome of LHR cAMP signalling is the synthesis of steroids hormones by gonads cells. The steroidogenesis machinery is controlled by PKA-dependent phosphorylations, and by LHR-dependent gene expression of several components of the machinery. Steroids finely regulate reproduction, and notably, under LHR activity, promote spermatogenesis in male and ovulation in female^12^. To assess the role of both LHR and Gαs trafficking in steroids synthesis, we used mouse Leydig cells with endogenous expression of mLHR and steroidogenic ability. It has been published that rodent LHR has slower internalization rate than the human receptor^45^, and it is yet unexplored whether it traffics to VEEs or to different endocytic compartments. Nevertheless, Lyga et al. have demonstrated that mLHR displays LH-induced persistent cAMP signalling in ovarian follicles, which occurs from intracellular sites^16^. Here, we show that hCG induces mLHR internalization in mLTC-1 cells (figure 5A). Importantly, inhibition of mLHR endocytosis by Dyngo4a or PitStop2 affected cAMP signalling dynamics in the same way as for the human receptor (figures 5B, 5C, S7), consistent with our finding in HEK293A cells. These data provide the first evidence that mLHR displays sustained Gαs/cAMP signalling from intracellular sites in male gonads cells. In addition to cAMP response within an hour, we show that 60-second ligand stimulation is enough to stimulate long-lasting cAMP and steroids accumulation (figure 5D, 5E). Notably, steroids accumulated from 60 minutes after ligand stimulation, suggesting steroidogenesis machinery in Leydig cells is already primed to rapidly respond to agonist stimulation (figure 5E). Preventing receptor internalization with Dyngo4a or PitStop2 strongly inhibited ligand-induced cAMP and steroids accumulation (figure 5D, 5E). Based on these results, we concluded that LHR endocytosis and intracellular Gαs/cAMP signalling are both necessary and sufficient to induce steroidogenesis. When performing continuous ligand stimulation for 3 hours, Dyngo4a only had partial effect on cAMP and steroids accumulation (figure 5F, 5G), though its effect was drastic even 6 hours after “pulse-chase” ligand stimulation (figure 5D, 5E). This suggested that under continuous exposure of cell surface receptors to ligand, Dyngo4a was permissive and allowed some receptor endocytosis in Leydig cells. The half maximal inhibitory concentration (IC50) of Dyngo4a on dynamin I and II being 0.38µM and 2.3µM, respectively^22^, it is possible that Dyngo4a, used at 30µM in our conditions, progressively lost efficacy over long period of time and continuous agonist stimulation. However, Dyngo4a still tended to reduce steroids accumulation, and PitStop2 maintained a strong inhibitory effect on progesterone accumulation (figure 5F, 5G). These observations were confirmed in mouse primary Leydig cells, in which both Dyngo4a and PitStop2 significantly inhibited hCG-induced testosterone production without drugs pre-incubation, showing that LHR endocytosis is required to produce steroids in primary gonads cells (figure 5I). This was supported by Gαs sequestration at the PM using Nb37-CAAX, which strongly inhibited progesterone production in mLTC-1 cells (figure 5H). Altogether, our results demonstrate that both LHR and Gαs endocytosis, that drive cAMP signalling from endosomes, are essential to steroidogenesis in Leydig cells.

Our study focuses on the functional relevance of LHR and Gαs trafficking and signalling from endosomes. Although the LHR mostly traffics to VEEs following endocytosis, a small fraction of receptors is internalized into EEs^13,15^. While it is unknown whether LHR can signal from EEs, the contribution of receptors in these distinct endosomal compartments for cAMP signalling is still do be determined. However, such study would require prior biochemical characterization of VEEs. It is also to be considered that Gαs protein exists in different isoforms, whose expression differs from distinct cell types and tissues^46,47^. Notably, the long and short variants of Gαs may not have similar functions^47^, and our experiments with transfection of exogenous Gαs were performed using short variants of Gαs. Nevertheless, using distinct cell types with endogenous Gαs expression, we confirmed the functional relevance of our observations. Besides G proteins, β-arrestins are important signalling partners of GPCRs, primarily known to mediate receptor internalization. Recent studies have put into light that not only some GPCRs can activate G proteins and signal from endosomes independently of β-arrestins, but can also undergo β-arrestin-independent internalization^48,49^. Regarding LHR, whether β-arrestins “enable”, “enhance” or “sculpt” Gαs/cAMP signalling from endosomes^48^ needs further investigation. Finally, we assessed the functional role of LHR internalization using chemical inhibitors of dynamin and clathrin, thus preventing not only LHR’s but many membrane proteins’ internalization. If available in the future, biased ligands able to specifically elicit LHR Gαs/cAMP signalling from PM or endosomes would be valuable tools to further decipher and confirm temporality, intensity and functional role of compartment-specific signalling.

In summary, this study provides evidence that LHR and Gαs trafficking are critical for cAMP signalling from endosomes, that drives nuclear cAMP accumulation, CRE-dependent gene expression and steroidogenesis. While relevance of GPCRs endocytosis is quite established, evidence on the role of Gαs translocation to endosomes to drive endosomal signalling and nuclear responses is emerging^24,25,36,43^. Our findings suggest that the role of Gαs translocation may be conserved across GPCRs, and shed light on the relevance of both receptor and Gαs trafficking at the physiological level. By demonstrating that LHR Gαs/cAMP signalling from endosomes drives steroidogenesis, this study opens new avenues for better understanding the etiology of reproductive pathologies such as polycystic ovary syndrome (PCOS), and for refining fertility control in ART and contraception^12,50–52^.

## Abbreviations and nomenclature

The abbreviations used are:

ART: assisted reproduction treatments;
AUC: area under the curve;
AVP: vasopressin;
BRET: Bioluminescence Resonance Energy Transfer;
β2AR: β2-adrenergic receptor;
cAMP: cyclic adenosine monophosphate;
CRE: cAMP responsive element;
CGRP: calcitonin gene-related peptide;
CRE: cAMP-responsive element;
DMSO: dimethylsulfoxide;
EE: early endosome;
ERK1/2: extracellular signal-regulated kinases 1/2;
FRET: Fluorescence Resonance Energy Transfer;
GPCR: G protein-coupled receptor;
GTP: guanosine triphosphate;
hCG: human chorionic gonadotropin;
HEK293A: human embryonic kidney 293A cells;
IC50: half maximal inhibitory concentration;
ISO: isoproterenol;
LH: luteinizing hormone;
LHR: luteinizing hormone receptor;
mLHR: mouse luteinizing hormone receptor;
mLTC-1: mouse Leydig Tumour Cells-1;
mNG: mNeonGreen;
NES: nucleus exclusion signal;
NLS: nuclear localization sequence;
OT: oxytocin;
PCOS: polycystic ovary syndrome;
PM: plasma membrane;
PTHR: parathyroid hormone receptor;
sAC: soluble adenylate cyclase;
TGN: trans-Golgi network;
VEE: very early endosome;
V2R: vasopressin type 2 receptor;
YFP: yellow fluorescent protein;
17βHSD3: 17β-hydroxydehydrogenase isoform 3.

## RESOURCE AVAILABILITY

### Lead contact

Additional information and request for resources or reagent should be directed to and fulfilled by the lead contact, Frédéric Jean-Alphonse.

### Materials availability

Data will be made available upon request to the lead contact. All materials generated in this study can be provided pending a completed material transfer agreement. Requests should be submitted to the lead contact Frédéric Jean-Alphonse.

## Acknowledgments

The authors thank Asuka Inoue (Tohoku University, Kyoto University, Japan) for kindly providing the HEK293ΔGs cells, Lucile Drobecq (CNRS, Nouzilly, France) for generating the LHR-LgBiT plasmid, Danièle Klett (CNRS, Nouzilly, France) for mLTC-1 cells, and the Merck company for kindly providing purified recombinant hCG (Ovitrelle^®^) and human LH (Luveris^®^). This work has benefited from the equipment and expertise of the Imaging facility “Plateau d’Imagerie Cellulaire” (PIC) (DOI: http://doi.org/10.17180/arap-gj59) and from the “Laboratoire Phénotypage-Endocrinology” (LPE) of UMR-PRC.

## Author contributions

F.J-A. conceptualization; J.G. and F.J-A. data curation; J.G. and F.J-A. formal analysis; F.J-A. and E.R. funding acquisition; J.G., M.H-H. and F.J-A. investigation; V.J., F.G. and F.J-A. methodology; E.R. and F.J-A. project administration; V.J., C.G. and S.A. resources; F.J-A. and E.R. supervision; J.G., E.R. and F.J-A. validation; J.G. and F.J-A. visualization; J.G. and F.J-A. writing-original draft; J.G., F.G., E.R. and F.J-A. writing-review and editing.

## Funding and additional information

This research was funded with the support of the Institut National de la Recherche pour l’Agriculture, l’Alimentation et l’Environnement (INRAE), Digit-Bio metaprogram (INRAE), the MAbImprove Labex (ANR-10-LABX-53), the Région Centre Val de Loire ARD2020 Biomédicaments SELMAT grant, the Bill & Melinda Gates Foundation CONTRABODY grant, and of the ANR-22-CE14-0050 MOSDER grant.

J.G. was funded by a joint fellowship from MabImprove Labex and Bill & Melinda Gates foundation. M.H-H. was funded by the Bill & Melinda Gates foundation. V.J. was funded by Région Centre Val de Loire ARD2020 Biomédicaments SELMAT grant. C.G., S.A., F.G. and E.R. were funded by INRAE. F.J-A. was funded by Centre National de la Recherche Scientifique (CNRS).

## 5. STAR Methods

### 5.1. Ligands and drugs

Recombinant human Chorionic Gonadotropin (hCG) (Ovitrelle^®^) was kindly provided by Merck and diluted in mQ H_2_O. AVP was purchase from Bachem (#4031337). Isoproterenol was purchase from Sigma-Aldrich (I6504). Dyngo4a was purchased at MedChemExpress (#HY-13863) and used at a final concentration of 30µM. PitStop2 was from Sigma-Aldrich (#SML1169) and used at 30µM.

The hCG-mNeonGreen (hCG-mNG) was produced in ExpiCHO-S^TM^ cells. ExpiCHO-S^TM^ cells at 6×10^6^ living cells/mL density were transfected with 20µg of pcDNA3.1 coding for the hCG-mNG, according to ExpiCHO-S^TM^ Expression System (Gibco^TM^) manufacturer’s instructions. Cells were cultured for one day at 37°C/8% CO_2_ on a shaker platform (120rpm) before supplementation with feed and enhancer solution (provided in the ExpiCHO-S^TM^ expression system transfection kit). Cells were then cultured for 12 days at 32°C/5% CO_2_ at 120rpm. The culture was centrifuged at 500g/30min/4°C, and the supernatant was then collected. The supernatant was then centrifuged again at 5000g/30min/4°C. The newly collected supernatant was dialyzed in regenerated cellulose membrane tubing with a 6-8kDa MWCO (Spectra/Por^M^) against a pH 8.0 buffer (50mM TrisHCl, 100mM NaCl) overnight at 4°C under continuous stirring. The dialyzed supernatant was centrifuged (5000g/30min/4°C) to remove insoluble debris, and passed through a buffer equilibrated (pH 8.0, 50mM TrisHCl, 100mM NaCl) Protino® Ni-IDA packed column (Macherey-Nagel^TM^). The column was washed with three times its volume of the following buffers: i) pH 8.0, 50mM TrisHCl, 100mM NaCl, ii) pH 8.0, 50mM TrisHCl, 1M NaCl, iii) pH 8.0, 50mM TrisHCl, 100mM NaCl. The hCG-mNG elution was performed with a pH 8.0, 50mM TrisHCl, 100mM NaCl, 500mM imidazole buffer. The hCG-mNG was buffer exchanged into a pH 8.0, 50mM TrisHCl, 100mM NaCl buffer, using 10DG Desalting Prepacked Gravity Flow Columns (BioRad Laboratories). The hCG-mNG was then concentrated using 30kDa MWCO Amicon Ultra centrifugation unit (Sigma-Aldrich) according to the manufacturer’s instructions. After analysis by Coomassie blue SDS-PAGE gel staining, purified hCG-mNG was stored at -20°C.

### 5.2. Cell culture

#### 5.2.1. Cell lines

Human Embryonic Kidney 293 (HEK293A) (Thermo Fisher Scientific, #R70507) and HEK293ΔGs (kind gift from A. Inoue) were cultured in DMEM (Eurobio, #CM1DME68-01). Mouse Leydig Tumour Cells 1 (mLTC-1) (ATCC, #CRL-2065) were cultured in RPMI 1640 (Eurobio, Cat #CM1RPM08-01). All mediums contained Glutabio and NaHCO_3_ and were supplemented with 10% heat inactivated fœtal bovine serum (Eurobio, #CVFSVF00-01), 100 IU/mL penicillin and 0.1 mg/mL streptomycin (Eurobio, #15140-122). Cells were kept at 37°C in a humidified 5% CO_2_ incubator. Cells were checked monthly for mycoplasma using the MycoAlert Mycoplasma Detection kit (Lonza, #LT07-318).

#### 5.2.2. Primary mouse Leydig cells

Primary mouse Leydig cells extraction and culture was conducted in L15 medium (Sigma-Aldrich, Cat #L4386) supplemented with 100IU/mL penicillin and 0.1mg/mL streptomycin (Eurobio, Cat #15140-122). All experiments were performed on animal materials post-mortem. Twenty testes from ten weeks-old C57BL/6JOlaHsd male mice (Inotiv, #057) euthanized by cervical dislocation were recovered. The albuginea was removed and testes were crushed with pliers, prior 15 minutes incubation in L15 medium containing 0.5mg/mL *Clostridium histolyticum* collagenase (Sigma-Aldrich, #C2674) and 0.01mg/mL bovine pancreas désoxyribonuclease I (Sigma-Aldrich, #Dn25), under 100rpm shaking in a 37°C water bath. After sedimentation of seminiferous tubules, supernatant was retrieved and centrifuged (218g, 8min, 15°C). The cell pellet was then resuspended in L15 medium and transferred to a 17%-42%-70% gradient of Percoll (Sigma-Aldrich, #P1644). After centrifugation (314g, 12min, 15°C), Leydig cells were collected at the 17%-42% interface. Leydig cells were washed with L15 medium, centrifuged (314g, 8min, 15°C), and the pellet was resuspended in L15 medium. Leydig cells were cultured the same day at a concentration of 3 to 4 million cells/mL, under 100rpm shaking in a 34°C water bath.

### 5.3. Plasmids

The cAMP BRET sensor NanoCUTie was designed based on the FRET sensor CUTie^27^ ; the YFP was replaced by a Venus and the CFP by a NLuc. DNA synthesis was done by Twist Bioscience. The nucleus addressed NanoCUTie was generated by subcloning NanoCUTie into pcDNA3.1 plasmid containing a Nuclear Localization Signal (NLS) addressing sequence in N-terminal position.

Nb37-mCherry was kindly given by Prof. J.-P. Vilardaga (University of Pittsburg, PA, USA). The plasma membrane addressed Nb37-mScarlet-I-(G4S)3x-CAAX (here after denoted Nb37-CAAX), was designed by adding the 13 last amino acids of Kras protein in C-terminal position, and replacing the mCherry by a mScarlet. The nucleus addressed Nb37-mScarlet-I-(G4S)3x-NLS (hereafter denoted Nb37-3xNLS) was designed similarly, by addition of 3 Nuclear Localization Signals sequences. Nb37-SmBiT was designed with a C-terminal (G4S)3x linker followed by the SmallBiT (SmBiT) sequence (VTGYRLFEEIL, Promega). All three cDNA were custom generated by Twist Bioscience.

The cAMP BRET sensor NLuc-Epac-VV ^20^ was kindly given by Prof. Kirill A. Martemyanov (The Scripps Research Institute Florida, FL, USA). The LHR plasmid was kindly given by Prof. Alfredo Ulloa-Aguirre (National University of Mexico, Mexico), whereas Flag-LHR was kindly given by Prof. Aylin Hanyaloglu (Imperial College London, United Kingdom). The mCherry plasmid was kindly given by Prof. Peter Friedman (University of Pittsburg, PA, USA). The pSOM-Luc plasmid was kindly given by Prof. Bernard Peers (University of Liège, Belgium). The Gαs-LgBiT, SmBiT-Gβ1, Gβ1, Gγ2, and Ric8B plasmids were kindly provided by Prof. Asuka Inoue (Tohoku University, Japan). The Gαs-ONE-GO plasmid was a gift from Prof. Mikel Garcia-Marcos (Addgene kit #1000000224). All other plasmids were generated by our group. Notably, Myr-Gs-YFP was designed by addition of myristoylation site in N-terminal position according to the construct by Prof. Wedegaertner’s group^53^. LHR-LgBiT were obtained by subcloning LHR-Rluc8 (kindly provided by Prof. Aylin Hanyaloglu, Imperial College London) with a pcDNA3.1 plasmid encoding either the (G4S)3x-LgBiT (from Dr. Lucie Pellissier) using the restriction sites NheI & XhoI. Plasmid encoding for pcDNA3.1-mTurquoise2-3xNLS was custom synthesized by Twist Bioscience.

### 5.4. Signalling and trafficking kinetics measurements by Bioluminescence Resonance Energy Transfer (BRET) and bioluminescence assays

Forty thousand HEK293A or sixty thousand mLTC-1 cells per well were seeded in 96-well plates (Greiner Bio-One 655098, 655083 or SPL #30196) previously treated with poly-lysine (Sigma-Aldrich, P8920) diluted at 0.01% in Ca^2+^-Mg^2+^-free PBS (Eurobio, #CS1PBS01-01), and then transiently transfected in suspension using Metafectene Pro transfection reagent (Biontex Laboratories, #T040-5.0) according to the manufacturer’s instructions, using the following DNA amounts:

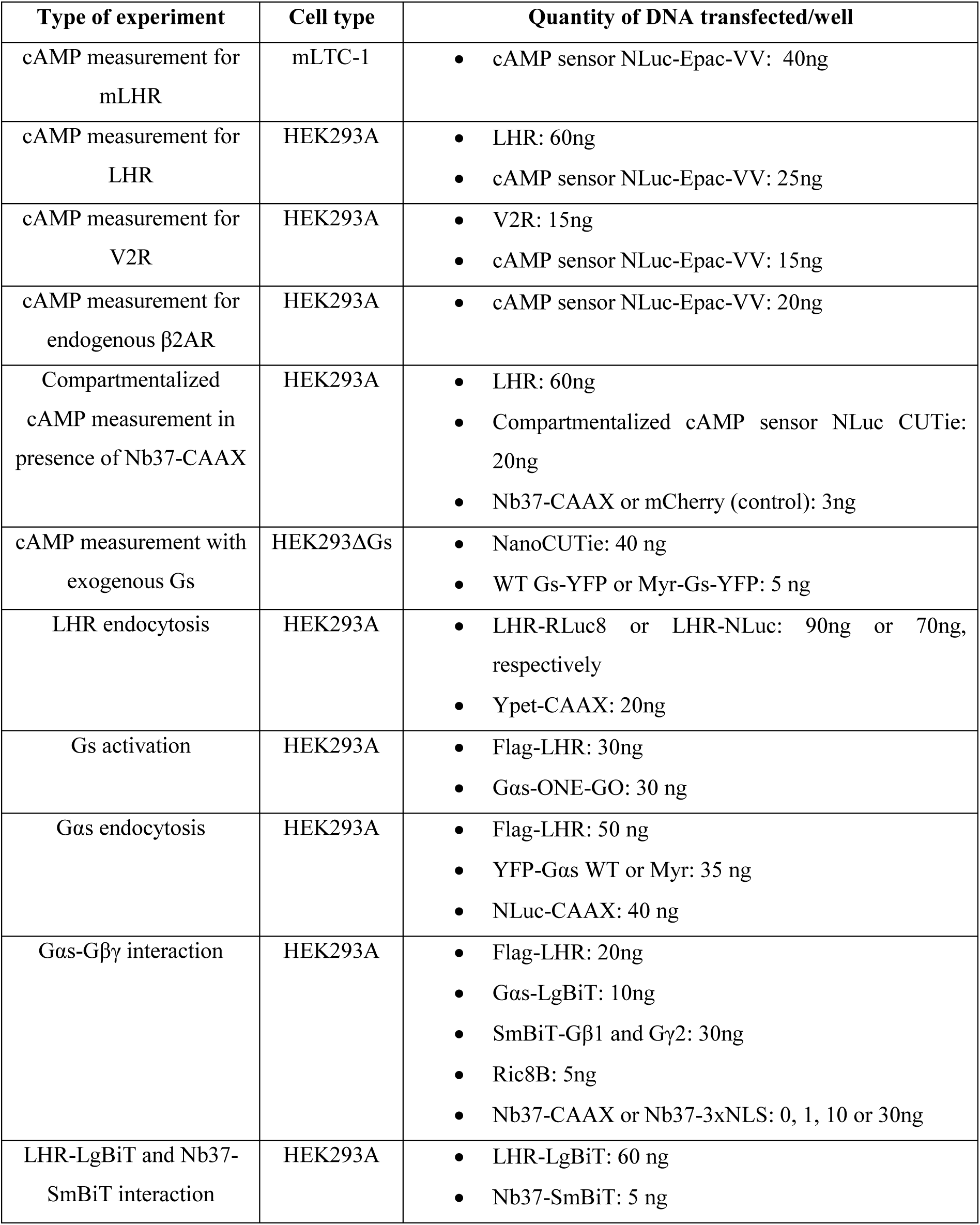

24 to 48 hours after transfection, BRET or bioluminescence measurements were performed upon addition of 5µM cœlenterazine H (Interchim, #R3078C) or 10µM furimazine (Promega) diluted in Ca^2+^/Mg^2+^-free PBS, containing no or different concentrations of hCG, and a drug for concerned conditions. For experiments performed in presence of Dyngo4a or PitStop2, cells were pre-incubated 30-35 minutes in presence of drug-containing buffers before measurements.

Signals were recorded for at least 30 minutes with a Mithras LB 943 plate reader (Berthold Technologies GmbH & Co., Wildbad, Germany). BRET ratios were calculated as follows: 480nm/540nm for NLuc-Epac-VV sensor experiments; 540nm/480nm for NanoCUTie sensors and receptor internalization experiments. Bioluminescence data were expressed as the percentage of control’s maximum.

### 5.5. cAMP accumulation measurement by HTRF

Sixty thousand mLTC-1 cells per well were seeded in 48-well plates 48 hours before the experiment, and were starved using RPMI medium without supplementation 17 hours before starting the experiment. Cells were pre-incubated for 35 minutes with either Dyngo4a or PitStop2 diluted in RPMI, or with RPMI supplemented with the equivalent amount of DMSO for control condition. Medium was removed, and cells were stimulated either transiently (60 seconds) or continuously with different concentrations (0.1 to 10 nM) of hCG diluted in RPMI containing either DMSO, Dyngo4a or PitStop2. After 3 hours, supernatants were collected and frozen at -20°C for further use. After supernatants dilution in RPMI medium, cAMP was quantified using cAMP Gs dynamic Kit (Revvity, Cat #62AM4PEB), following the manufacturer’s instructions. Fluorescence measurement was performed by donor excitation at 620nm and acceptor emission reading at 665nm, using a TriStar^2^ LB 942 Multimode Microplate Reader (Berthold Technologies GmbH & Co.). The 665nm/620nm ratio was calculated, and cAMP concentrations were determined using standards, following the manufacturer’s instructions.

### 5.6. CRE-dependent gene transcription quantification

Forty thousand HEK293A cells per well were seeded in 96-well plates previously treated with poly-lysine (Sigma-Aldrich, Cat #P8920) diluted at 0.01% in Ca^2+^-Mg^2+^-free PBS (Eurobio, Cat #CS1PBS01-01), and then transiently transfected in suspension using Metafectene Pro transfection reagent (Biontex Laboratories, Cat #T040-5.0) according to the manufacturer’s instructions, using the following DNA amounts: 60ng/well Flag-LHR, 40ng/well pSOM-Luc, plus 30ng/well of Nb37-CAAX or mCherry for concerned experiments. The pSOM-Luc plasmid consists in the firefly luciferase reporter gene under the control of somatostatin promoter region cAMP responsive element (CRE), and allows to assess CRE-dependent gene transcription. After 48 hours transfection, cells were pre-incubated for 35 minutes with DMEM containing Dyngo4a, PitStop2 or equivalent volume of DMSO (control condition), for experiments with chemical inhibitors. No pre-incubation was performed for experiments with Nb37. Medium was then removed, and cells were stimulated during 6 hours with hCG (concentrations from 0.01 to 10nM) diluted in DMEM. For experiments with chemical inhibitors, hCG was diluted in DMEM with either DMSO, Dyngo4a or PitStop2. Supernatants were removed and cell lysis was induced by adding Bright-Glo Luciferase assay substrate (Promega, Cat #E2620). After 5 minutes incubation at room temperature and away from light, luminescence was quantified with a Mithras LB 943 plate reader (Berthold Technologies GmbH & Co.).

### 5.7. Imaging

HEK293A and mLTC-1 cells were seeded on 35 mm glass bottom dishes (Greiner BIO-One # 627870 and #627860) previously coated with collagen (Gibco, #A10483-01). After 6 hours, cells were transfected with plasmids encoding for the LHR and fluorescently-tag proteins using lipofectamine 3000 (Thermo Fisher Scientific) for 24 to 48 hours. For live cell imaging, cells were incubated in DMEM phenol red-free (Gibco, ref 21063-029) supplemented with 1% FBS, and images acquired by confocal microscopy (Zeiss LSM900-Airyscan2) with controlled temperature at 37°C. For pulse-chase experiments, hCG-mNG was added 60 seconds (HEK293A) or 120 seconds (mLTC-1) before being washed out twice with DMEM phenol red-free supplemented with 1% FBS.

### 5.8. Progesterone measurements

For experiments performed in presence of chemical internalization inhibitors, supernatants were obtained as described earlier (see 5.5). For experiments performed in presence of Nb37-CAAX, sixty thousand mLTC-1 cells per well were seeded in 48-well plates 48 hours before the experiment and were transiently transfected in suspension with 80ng per well of either mCherry or Nb37-CAAX, using Metafectene Pro transfection reagent (Biontex Laboratories, Cat #T040-5.0) according to the manufacturer’s instructions. mLTC-1 were starved using RPMI medium without supplementation 17 hours before starting the experiment. Medium was removed, and cells were stimulated with different concentrations (0.1 to 10 nM) of hCG diluted in RPMI. After 3 hours, supernatants were collected and frozen at -20°C for further use. Progesterone concentration in each supernatant was determined using enzyme-linked immunoassay (EIA) adapted from Canepa et al.^54^. Briefly, 96-well plates were coated with 0.2µg/well of anti-mouse IgG goat IgG at 4°C for at least one night. After three washes (25mM Tris, 37mM NaCl, 0.5mM MgCl2, pH 7.5, 0.05% v/v Tween 20), plates were saturated for 4 hours with Tris/BSA buffer, before overnight incubation at 4°C with anti-progesterone mouse monoclonal antibody and 25µL of each sample or standard. Standards at a concentration of 0.25 to 32ng/mL progesterone (Steraloid, Cat #Q2600) were prepared in Tris/BSA buffer. Sample supernatants were diluted at 1/75 in the Tris/BSA buffer. After incubation, a solution of progesterone alkaline phosphatase was added prior 1-hour incubation under constant shaking away from light. After washes, a solution of p-nitrophenyl phosphate (Biorad, Cat #BUF044B) was added in each well. A 2-hours incubation at 37°C was conducted, and absorbance was quantified at 405nm with a microplate reader (Tecan). The sensitivity of the assay was 0.4ng.mL^-1^, and the inter-assay coefficient of variation was <6.7%. Progesterone concentration in samples supernatants was determined based on progesterone standards.

### 5.9. Testosterone measurements

Primary mouse Leydig cells at a concentration of 3 to 4 million cells/mL were stimulated with different concentrations of hCG diluted in L15 medium containing DMSO, 30µM Dyngo4a or 30µM PitStop2. After 3 hours, cells were centrifuged (200g, 5min) and supernatants were collected and frozen at -20°C for further use. Testosterone concentration in each supernatant was determined using a previously developed enzyme-linked immunoassay^55^.

### 5.10. Data normalization and statistical analysis

Data were obtained from at least three independent experiments (N), each containing two to three biological replicates (n). For BRET data, a normalization on baseline values was done to represent agonist-induced signals. Areas under the curve (AUC) were calculated for each independent experiment. Because of inter-assay signals variation, gene expression bioluminescence data and testosterone quantification in primary Leydig cells were expressed as the percentage of the control condition’s maximum. BRET and luminescence data were analysed and plotted using GraphPad Prism 10 (GraphPad Software Inc.). Values were represented as means ± SEM. Imaging data were analysed using the Fiji software. Statistical analysis was performed using GraphPad Prism 10 (GraphPad Software Inc.). Statistical significance was assessed by one-tailed Mann-Whitney test, to compare a given condition to the corresponding control. p-value was considered significant when ≤0.05, and indicated as follows: *p≤0.05, **p<0.01; ***p<0.001, ****p<0.0001. A statistical tendency (0.05 < p-value < 0.07) was indicated by #.

