## Supplementary Information for "LHR and Gαs trafficking drive sustained cAMP signalling from endosomes to control steroidogenesis"

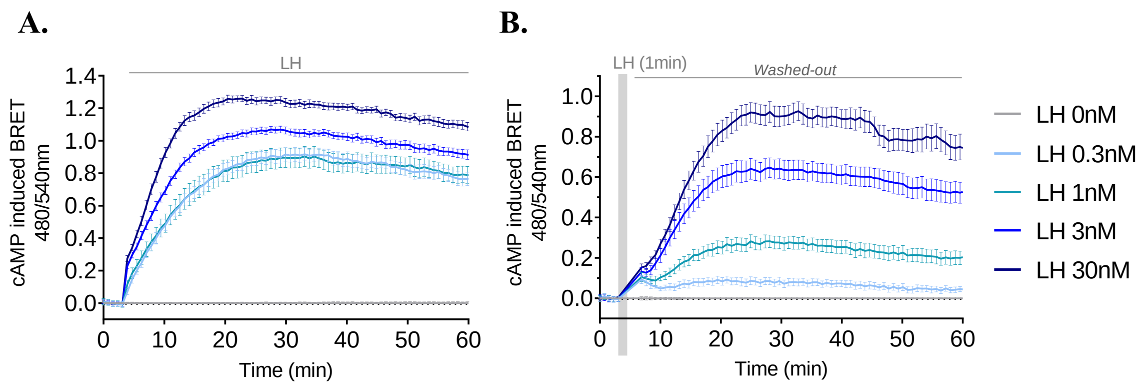


**<u>Figure S1</u>: LH-dependent LHR activation generates sustained cAMP signalling.** LHR-dependent time courses of cAMP signalling induced by continuous **(A)** or 60-second **(B)** LH stimulation of HEK293A cells transiently expressing the receptor. **N=3 (n=9)**, means ± SEM.

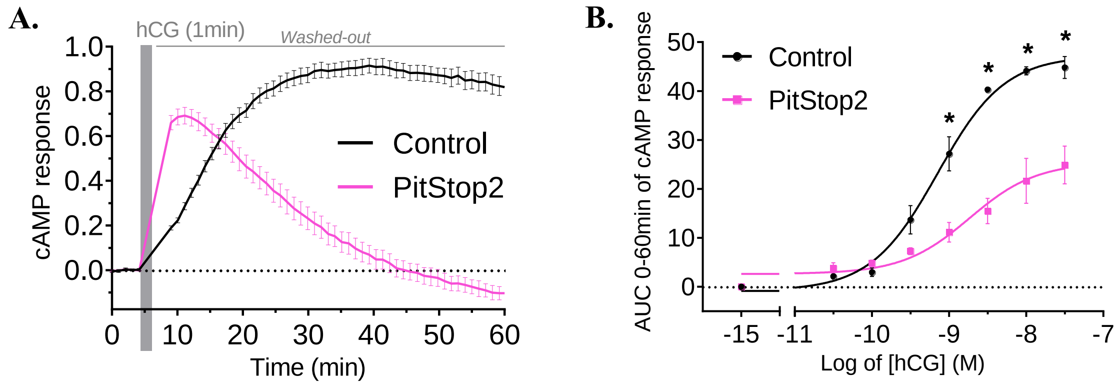


**<u>Figure S2</u>: PitStop2 inhibits endosomal cAMP signalling. (A)** Time courses of cytosolic LHR-dependent cAMP production dynamics in HEK293A cells, induced by 60-second stimulation with hCG, in presence or absence of 30 µM clathrin inhibitor PitStop2. **N=3 (n=9)**, means ± SEM. **(B)** Dose-response curves from area under the curve of cAMP response measured between 0 and 60 minutes, with or without 30µM PitStop2. **N=3**, means ± SEM.

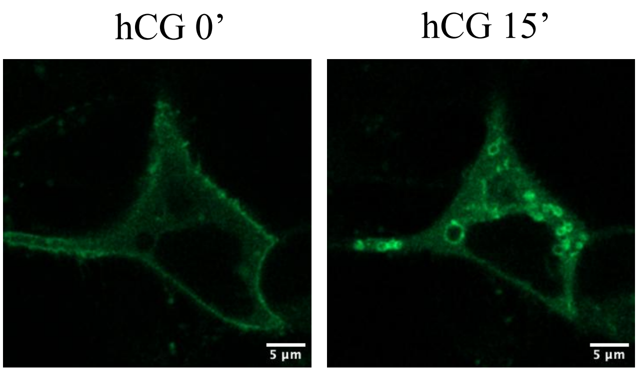


**Figure S3: Gαs endocytosis is dynamin-independent.** Gαs-YFP cellular localization in HEK293A cells expressing LHR, before (left) or after (right) 15 minutes of 30 nM hCG stimulation in presence of 30µM Dyngo4a. Scale bar = 10 µm.

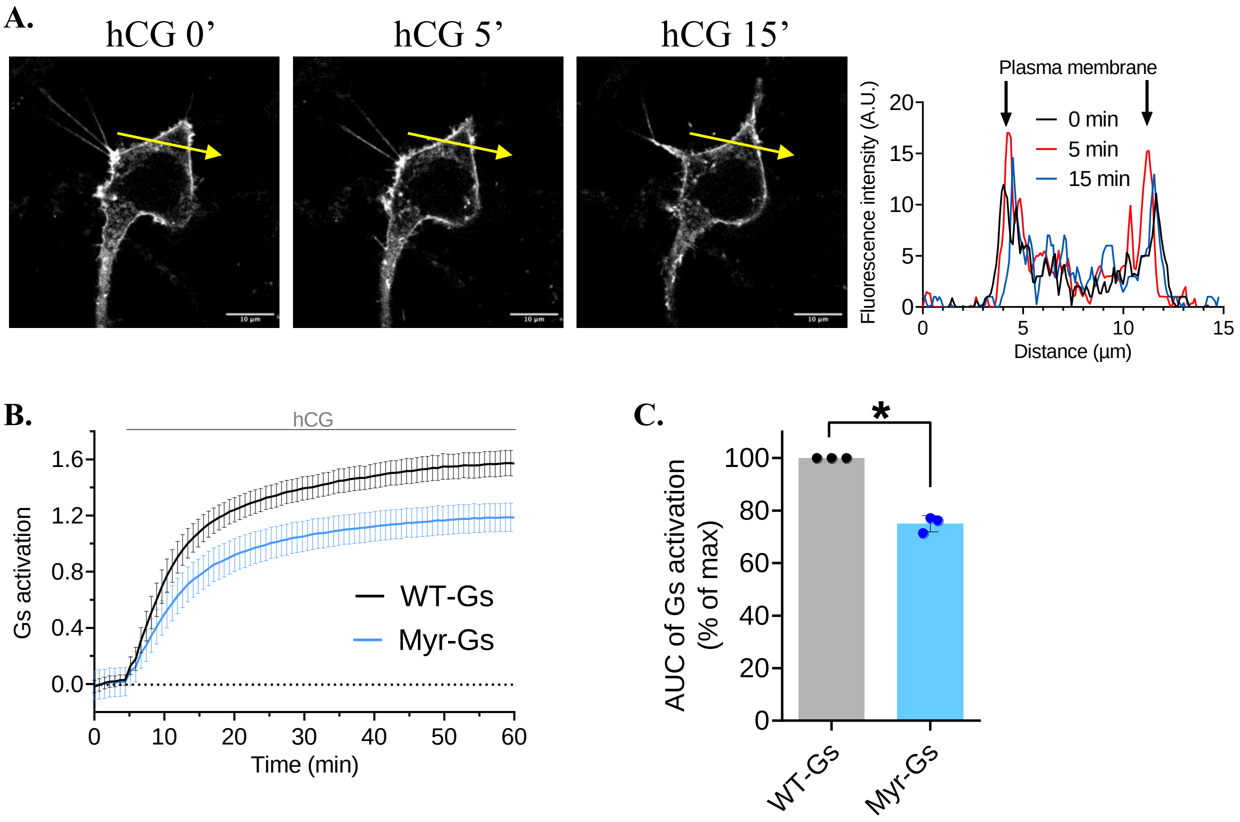


**Figure S4: Myr-Gs is localized at the plasma membrane**. **(A)** Imaging of Myr-Gs-YFP upon 30 nM hCG stimulation. The right panel indicates the fluorescence intensities from the arrow across time. **(B)** Luminescence complementation assay monitoring the kinetics of recruitment of Nb37-SmBiT to LHR-LgBiT by activated WT-Gs or Myr-Gs expressed in HEK293ΔGs cells. **N=3 (n=9)**, means ± SEM. **(C)** Area under the curve from the kinetics measured between 0 and 60 minutes. **N=3**, means ± SEM.

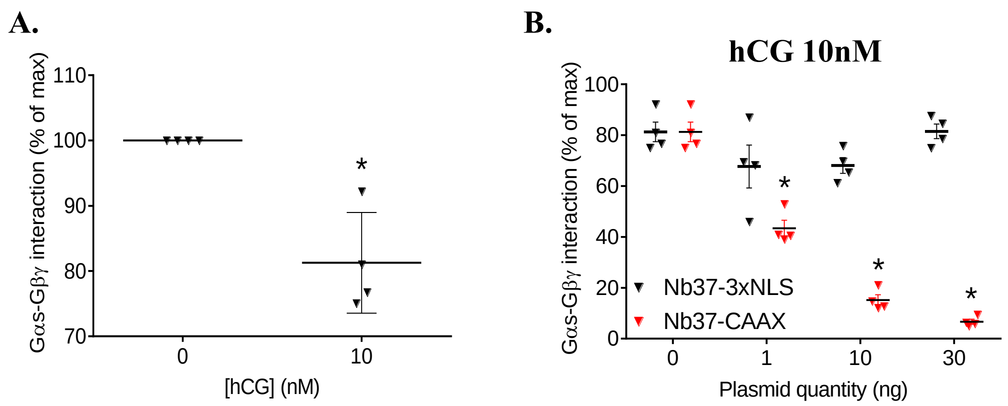


**<u>Figure S5</u>:** Inhibition of Gs activation by Nb37-CAAX. **(A)** Luminescence complementation assay monitoring the Gαs and Gβγ ligand-induced dissociation stimulated by 10 nM hCG. Data represent area under the curve of Gαs-LgBiT and Gβ1-SmBiT interaction between 0 and 10 minutes. Data were expressed as the percentage of control's maximum. N=4 independent experiments performed in triplicates, means ± SEM. **(B)** Effect of Nb37-3xNLS (nucleus localized) or Nb37-CAAX (plasma membrane localized) expression on Gαs-Gβγ dissociation stimulated by 10 nM hCG. Data represent area under the curve of Gαs-LgBiT and Gβ1-SmBiT interaction between 0 and 10 minutes. Data were expressed as the percentage of control's maximum. **N=4** independent experiments performed in triplicates, means ± SEM.

**A.**

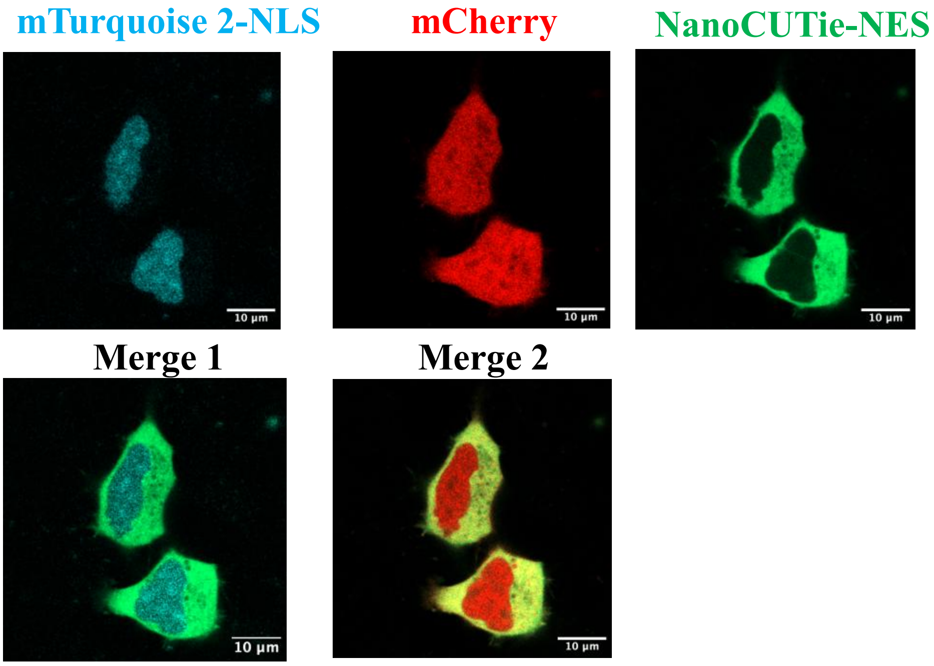


**B.**

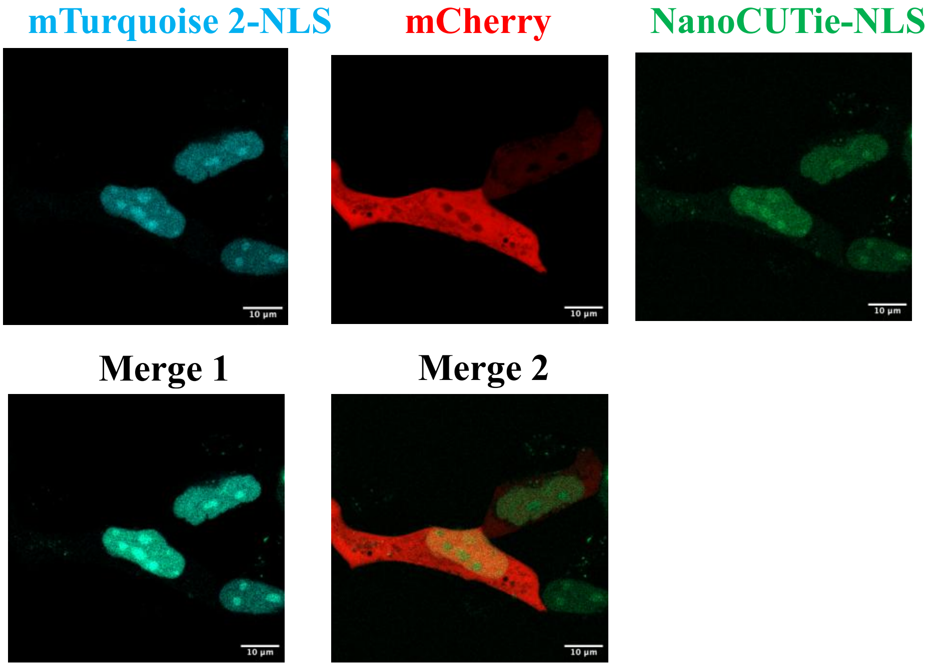


<u>**Figure S6**</u>**: Cellular localization of NanoCUTie sensor**. **(A)** Cellular localization of cytosol-addressed NanoCUTie (NanoCUTie-NES). Cellular localization of mTurquoise2 fused to a nuclear-localization sequence (NLS) and of mCherry without addressing sequence were used as references. Merge 1: mTurquoise2-NLS and NanoCUTie-NES channels. Merge 2: mCherry and NanoCUTie-NES channels. **(B)** Cellular localization of nucleus-addressed NanoCUTie (NanoCUTie-NLS). Cellular localization of mTurquoise2 fused to a nuclear-localization sequence (NLS) and of mCherry without addressing sequence were used as references. Merge 1: mTurquoise2-NLS and NanoCUTie-NLS channels. Merge 2: mCherry and NanoCUTie-NLS channels. Scale bar = 10µm.

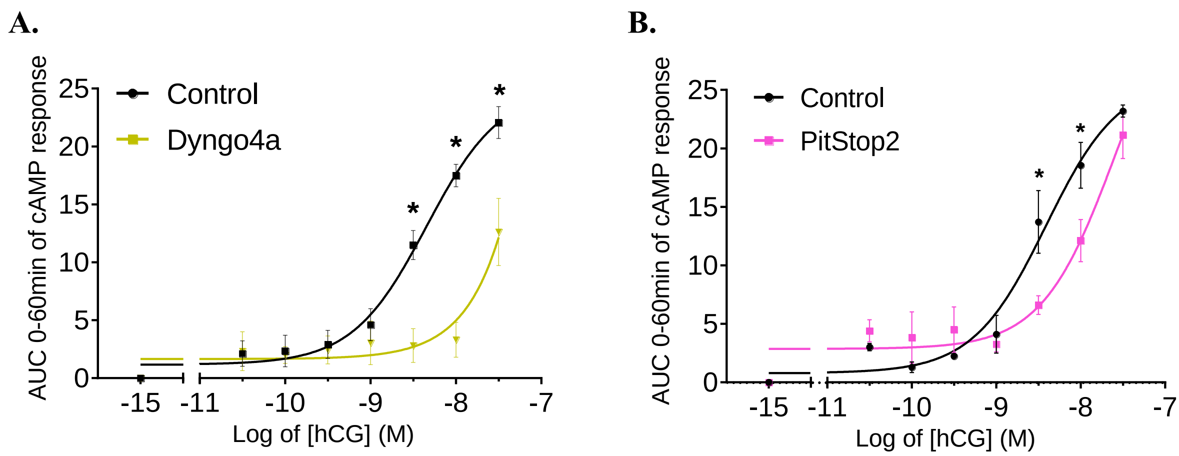


**<u>Figure S7</u>: mLHR displays sustained cAMP signalling that requires receptor internalization**. Representation at each hCG concentration of the area under the curve from cAMP response measured in mLTC-1 cells between 0 and 60 minutes, with or without 30µM Dyngo4a (A) or Pitstop2 (B). **N=3**, means ± SEM.
